# Limitations of *p*_50_ as a measure of seed longevity and the way forward

**DOI:** 10.1101/2025.06.11.659081

**Authors:** Lea Klepka, Angelino Carta, Fiona R. Hay, Anna Bucharova

**Author notes:** Corresponding author: Anna Bucharova.

## Abstract

**Premise:** Comparative studies of seed longevity assess how environmental conditions, species identity, populations, or genotypes affect seed viability loss in stored seeds. These studies commonly use the time it takes for seed viability to drop to 50% - known as *p*_50_, - as a measure of seed longevity. However, *p*_50_ is influenced by the initial seed viability. To allow comparisons regardless of the initial seed viability, standard protocols for comparative studies of seed longevity recommend using seed lots with “similar and high” initial viability, typically between 85 and 100%. However, even such range of initial seed viabilities might result in substantial variation in p_50_.

**Methods:** Here, we use a modelling approach to illustrate how variation in initial viability affects *p*_50_ estimates, and to propose alternative, more robust measures of seed longevity in storage.

**Results:** We show that for hypothetical accessions with identical rates of seed viability loss, variation in initial viability between 85 and 100% leads to a threefold variation in *p*_50_ estimates. Most of the unintended variation is introduced by accessions with very high viability (>95%). Restricting the initial viability to a narrower range (e.g., 85–95%) reduces but does not eliminate this bias. Alternatively, *p*_50_ can be recalculated to a standardized value of initial viability (e.g., 90%), which makes it proportional to the rate of probit viability loss without any noise. However, the most straightforward measure of seed longevity for comparative studies is the rate of viability loss itself, represented by σ (sigma) from the viability equation.

**Conclusion:** The common measure of seed longevity, p50, is affected by unintended, substantial variation caused by varying initial viability of the seed lot, and is thus suboptimal for comparative studies of seed longevity among seed accessions. More robust measures of seed longevity include the rate of viability loss, or p_50_ standardized to a certain initial seed viability.

## Introduction

Seed longevity - the period over which seeds remain viable (Nadarajan et al., 2023) - is a critical seed trait. A detailed, comprehensive understanding of how long seeds maintain viability could facilitate effective seed storage in conservation seed banks and gene banks, and enhance understanding of seed persistence in the soil. There is a common understanding that seed longevity strongly depends on environmental conditions (Solberg et al., 2020; Corbineau, 2024) and varies between species (Probert et al., 2009; Merritt et al., 2014), but there is also some variation between accessions of the same species, both for crops and wild species (e.g., Probert et al., 2009; Mondoni et al., 2011; Lee et al., 2019; White et al., 2023; Balasupramaniyam et al., 2025).

Comparative studies of seed longevity among seed accessions typically expose seeds to specific constant environmental conditions, depending on the research questions, with seed survival then monitored at predefined intervals (e.g., Walters et al., 2005; Probert et al., 2009; Carta et al., 2018; Moravcová et al., 2022). Studies focusing on longevity of seeds in conservation seed banks commonly follow the comparative longevity protocol developed by the Millennium Seed Bank of the Royal Botanic Gardens Kew (Newton et al., 2014; Royal Botanic Gardens, Kew, 2022). Briefly, seeds are exposed to 60% relative humidity and 45 or 60°C, conditions that are assumed to accelerate chemical processes similar to the ones that happen in the seeds during long-term storage in ex situ facilities.

Individual samples are removed from the artificial storage conditions after 1, 2, 5, 9, 20, 30, 50, 75, 100 and 125 days, and the proportion of viable seeds is determined. The proportion of viable seeds (as a binomial variable) is then related to the time in aging conditions using a probit link function. The probit transformation converts binomial responses onto a linear scale with a normal distribution to fit the seed viability equation (Ellis and Roberts, 1980):

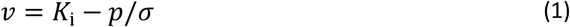

Where v is seed viability (in probits) of the seed sample after p days in the aging environment. *K*_i_ is the y-intercept, that is the theoretical initial viability (in probits), and σ (sigma) is the time for viability to fall by 1 probit. The σ parameter directly describes the rate of seed viability decay on a probit scale. To interpret *K*_i_ on a more intuitive scale it can be back-transformed from probits to percentages, *K*_i_(%), using the inverse of the probit function. *K*_i_, as the theoretical initial viability of a seed lot, estimated based on fitting the survival curve through probit analysis, is similar but likely not identical with the experimentally observed seed viability at the start of the aging experiment.

As a measure of seed longevity, comparative studies commonly use the time when germination upon removal from the aging environment has fallen to 50% of the tested seeds, the *p*_50._ (Priestley et al., 1985; Walters et al., 2005; Probert et al., 2009; Mondoni et al., 2011; Satyanti et al., 2018). However, *p*_50_ inherently depends on the initial viability, *K*_i_:

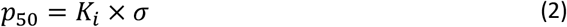

When the initial viability is higher, e.g., 95%, it takes longer for seed viability to fall to 50% compared to when the initial viability is lower, e.g., 85 or 70% (Figure 1). The initial viability *K*_i_ is a seed lot (accession) trait and can be affected by the age of the seeds at the beginning of the experiment, post-harvest treatments, or other environmental effects. Thus, any accession of any species could have any reasonable level of initial viability. While the initial viability is an important measure of seed quality, comparative studies are rather interested in the rate of seed viability loss, which is specific for species, populations or genotypes. This rate of seed viability loss is best represented by σ, which is independent of the initial viability. Consequently, Hay et al. (2022) state that σ would be the most suitable measure of seed longevity in storage. At the same time, the authors acknowledge that σ is not intuitive to understand, and recommend using *p*_50_ if all included accessions have similar and high initial seed viability. In that case, *p*_50_ should be proportional to σ (Hay et al., 2019, 2022).

**Figure 1.**
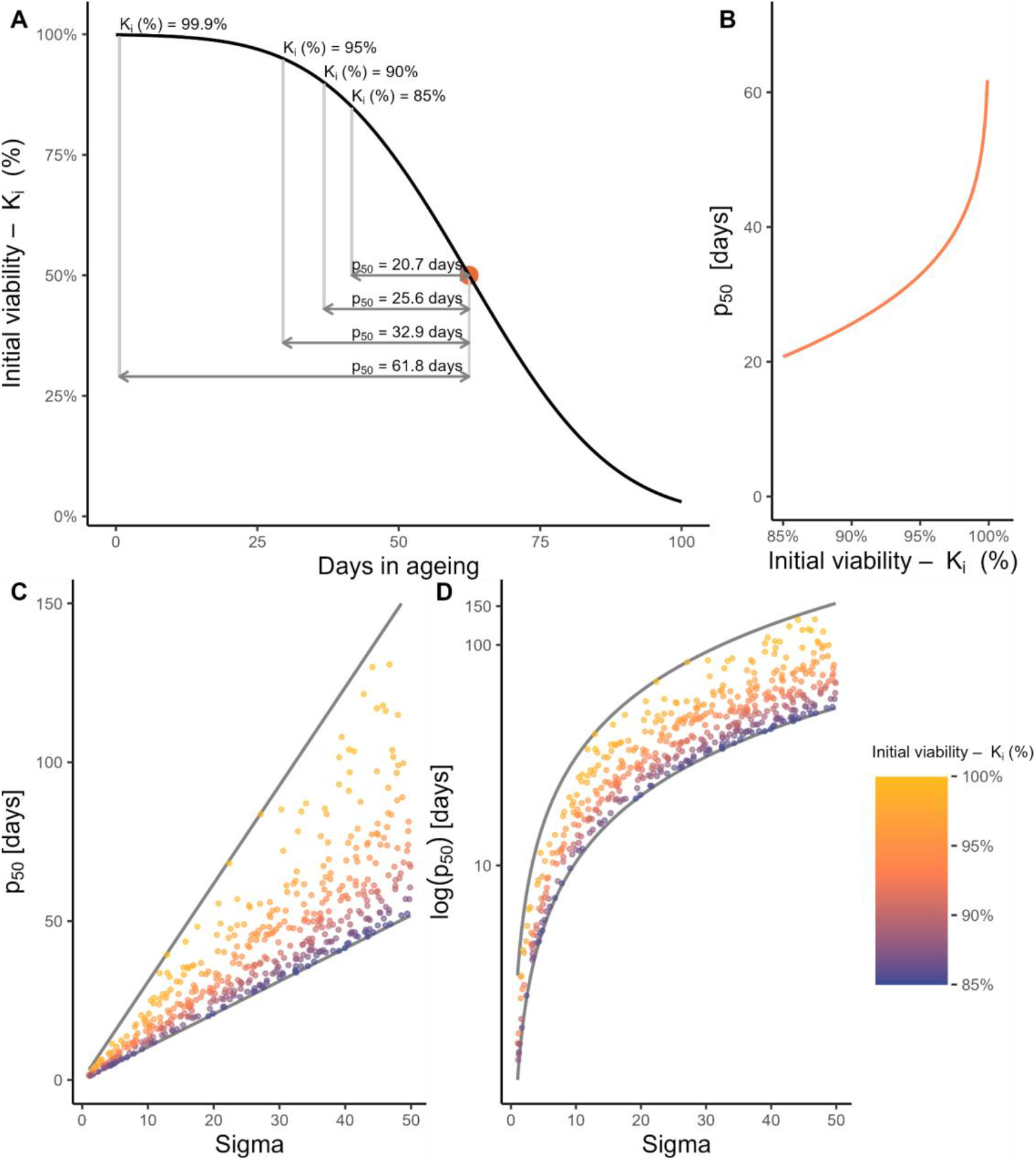
Illustration of how *p*_50_ estimates depend on the initial viability of a seed lot. (A) Seed survival curve describing the decline in viability of hypothetical seed accessions with a rate of probit viability loss, σ=20 days. (B) *p*_50_ exponentially increases as initial viability of the seed lot at the start of the aging experiment increases, here illustrated for a hypothetical accession with σ=20 days. (C) Illustration of the noise introduced to *p*_50_ estimates by using seed accessions with initial viabilities varying between 85 and 99.9%. Points are *p*_50_ estimates for 500 hypothetical seed accessions with σ randomly varying between 1 and 50 days and initial viabilities between 85 and 99.9%. (D) Is the same as C, with a logarithmic scale for the y-scale. The grey lines in C and D correspond to *p*_50_ with *K*_i_ fixed to 85 and 99.9%.

So far, the “similar and high initial viability” has not been clearly defined. In practice, most studies have used seeds with initial viability of at least 80% (Probert et al., 2009; Mondoni et al., 2011; Merritt et al., 2014; but see also Satyanti et al., (2018) who used 75%) while the protocol developed by the Millennium Seed Bank of the Royal Botanical Gardens Kew recommends using seeds with at least 85% initial viability (Royal Botanic Gardens, Kew, 2022). This allows for using seeds with initial viabilities ranging between 85 and 100%, which inevitably introduces some unintended variation to the *p*_50_ estimates, although in reality, a narrower range is likely covered, especially for prestored seed bank accessions. The question is whether the magnitude of this unintended variation is acceptable for comparative studies among accessions.

In this study, we illustrate the magnitude of the unintended variability in *p*_50_ caused by using seed lots that range in initial viability between 85 and 100%. While the results presented below might be obvious for seed physiologists familiar with probit analysis, they were unexpected to the first and last authors of this study - ecologists who only recently started to work on seed longevity. Given that *p*_50_ estimates from seed lots with varying initial viability are commonly used in comparative studies, we believe that illustrating the magnitude of the variability in *p*_50_ will be useful for fully comprehending the consequences of this practice. We then outline three possible solutions for how to reduce or remove the unintended variation in the seed longevity estimates.

## Methods

All simulations were performed in R (R version 4.4.2 (2024-10-31 ucrt)). As many biologists have a negative attitude to mathematical equations (Fawcett and Higginson, 2012), we do not provide equations in the main text but see e.g., Hay et al. (2014). We provide an annotated R-script in the supplementary materials (Supplementary Material 1), allowing readers to follow each calculation step of the models, replicate the results, and adjust thresholds to suit individual needs.

To achieve our goal, we adopted a three-step procedure: (1) we first illustrated the basic relationship between *p*_50_ and initial viability, *K*_i_ (%), then, (2) we illustrated the variation in *p*_50_ estimates across seed accessions that vary in initial seed viability. Finally, (3) we explored possible solutions to reduce or remove the unintended variation in *p*_50_ estimates.

In the first step, we illustrated the basic relationship between *p*_50_ and initial viability, *K*_i_(%), using a hypothetical accession with a fixed rate of probit viability loss, σ=20 days. This means that seed viability decreases by one probit – for example, from 84.1 to 50%, or from 97.7 to 84.1 % – every 20 days. While we kept σ constant, we varied initial seed viability in 0.1% steps between 85 and 99.9% (100% is theoretically impossible). We then calculated *p*_50_ using equation (2) and plotted it against initial seed viability.

In the second step, we illustrated the magnitude of unintended variation in *p*_50_ estimates that is introduced by using seed accessions that vary in initial seed viability. For this, we simulated 500 hypothetical seed accessions with σ randomly varying between 1 and 50 days and initial viability between 85 and 99.9%. This example mimics a comparative analysis across many accessions. We calculated *p*_50_ using equation (2) and then related the obtained *p*_50_ to the σ of the given accession. We further show the relationship between log(*p*_50_) and σ, as *p*_50_ is often log-transformed in comparative analyses (Probert et al., 2009; Merritt et al., 2014; Satyanti et al., 2018).

In the third step, we explored possible solutions to reduce or remove the unintended variation introduced to *p*_50_ estimates by using seed lots varying in initial viability. As one option, we explored the effect of restricting the range of initial viabilities to 75-85%, 80-90%, 85-95% or 90-99.9%. To do this, we first simulated 500 hypothetical seed accessions as in the paragraph above, with σ randomly varying between 1 and 50 days and initial viability within the given intervals. As an alternative option, we directly used the slope of probit viability loss, σ, although it might be hard to interpret. To help the reader to more intuitively understand the σ, we visualize the loss of viability that happens in σ days, depending on the initial viability. The last option we explored was to recalculate *p*_50_ based on a standardized value of *K*_i_, e.g., 90% (i.e., as a multiple of σ), so that it is independent of the true initial viability. We then report the *p*_50_ estimate as the time needed for viability of a seed lot to drop from 90 to 50%.

## Results

For a hypothetical seed accession with σ=20 days, the estimates of *p*_50_ varied between 20.7 and 61.8 days when calculated for seed accessions with initial viabilities between 85 and 99.9%, respectively.

The differences between *p*_50_ estimates were especially large when using seed lots with very high viabilities (Figure 1A, B). Independent of the σ value, the *p*_50_ based on a seed lot with 99.9% initial viability was 2.98-times larger than the *p*_50_ based on a seed lot with 85% initial seed viability, which introduces substantial noise to comparisons across accessions (Figure 1B). While log-transformation of the *p*_50_ reduced the noise to some extent, *p*_50_ based on a seed lot with 99.9% viability was consistently larger than *p*_50_ based on a seed lot with 85% viability, with an additive difference of 1.1 units on the logarithmic scale (Figure 1C).

Restricting the initial seed viability to a specific range also reduced the noise in the resulting *p*_50_. For example, when we restricted the initial viability to 85-95%, the difference between the highest and lowest possible *p*_50_ at the same σ was only 0.61 σ (i.e., the probit difference between 85 and 95%). In contrast, using seeds with initial viabilities ranging between 90-99.9%, this difference increased to 1.81 σ – almost three times larger (Figure 2). When we fixed the initial seed viability to 90%, or any other value, the *p*_50_ estimate increased proportionally with σ without any noise (Figure 2A).

**Figure 2.**
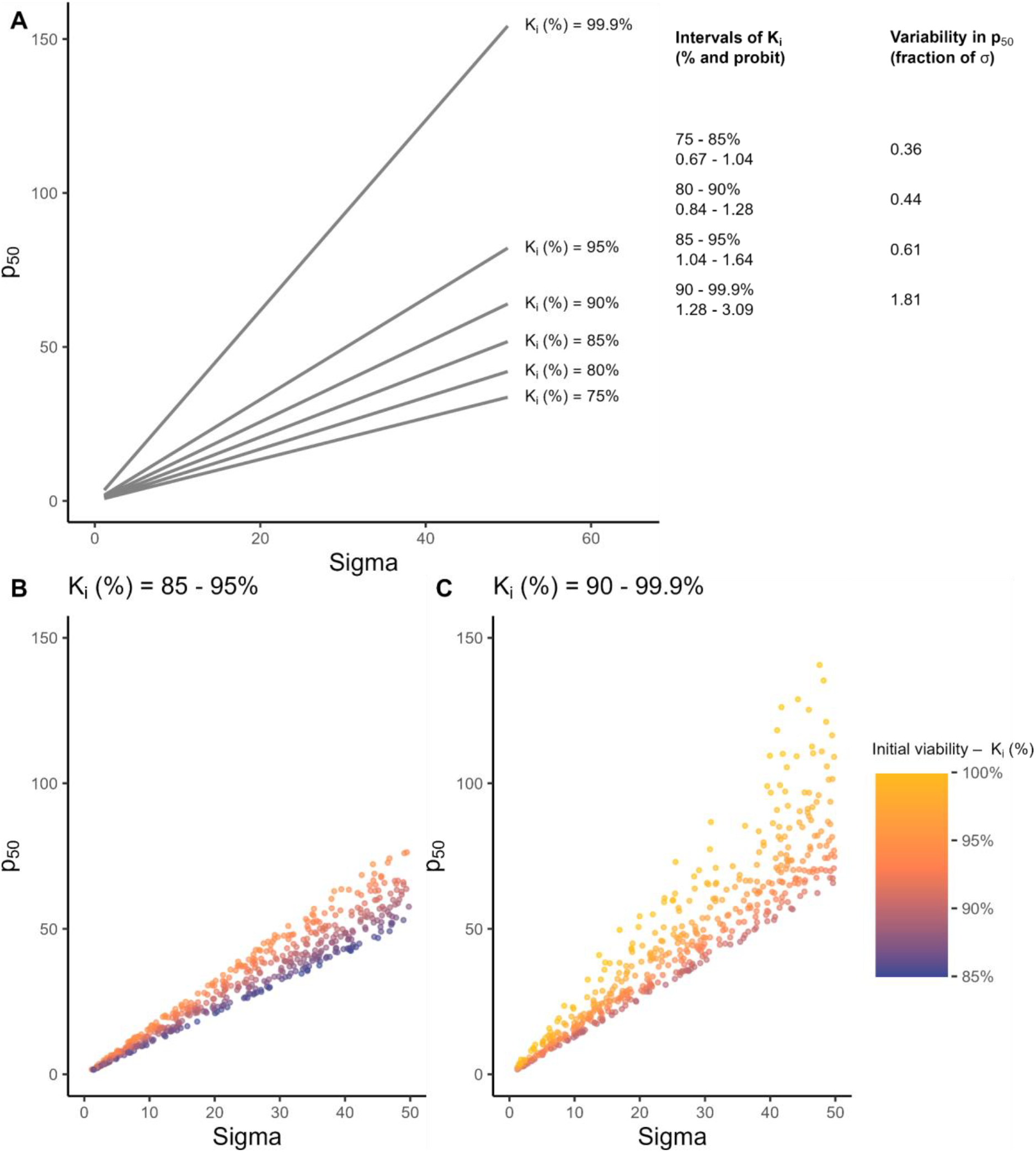
(A) Relationship between *p*_50_ and σ when the initial viability is fixed to a specific value. Note the absence of any variation in *p*_50_. On the right are variations in *p*_50_ estimates when only seed lots from a restricted range of initial viabilities are used. (B) and (C) illustrate the variation in *p*_50_ estimates for 300 hypothetical seed accessions that randomly vary in σ from 1 to 50 days (x-axis) and initial viability 85 to 95% (B) or 90 to 99.9 % (C).

## Discussion

Comparative studies of seed deterioration in storage typically involve exposing seeds to high temperature and humidity for different time periods to accelerate seed deterioration. As a measure of seed longevity, most studies use the *p*_50_ estimate, which is the time it takes for the viability of the seed lot to fall to 50% (i.e., half the seeds are able to germinate and half are not able to germinate and presumed dead). The *p*_50_ values are typically estimated using seed lots with high initial viability. However, here we illustrate that using seed lots with >85-100% initial seed viability introduces a three-fold difference in the resulting *p*_50_ estimates. To address this issue, we propose three potential solutions and alternatives to the *p*_50_ as currently in practice.

Most of the existing large comparative studies on seed longevity in storage have used *p*_50_ estimates based on seed lots with varying initial viability (e.g., Probert et al., 2009; Mondoni et al., 2011; Merritt et al., 2014; Satyanti et al., 2018). These *p*_50_ estimates are likely influenced by substantial variation in initial viability. While using seeds with higher initial viability improves the precision of the individual *p*_50_ estimates, here we show that including seed lots with 95-99.9% viability actually produces the largest variation in *p*_50_. Thus, the *p*_50_ estimates in these studies may mask subtler relationships between longevity and other factors, such as seed weight or climatic parameters. Log-transformation of the *p*_50_ estimates before relating it to environmental parameters transformed the noise from multiplicative to linear along the *p*_50_ values, but the data are still noisy and the impact of the noise on the results remains unclear.

The *p*_50_ estimates are calculated from the transformed seed viability curve, specifically from the intercept (theoretical initial viability) and slope, that is the rate of viability loss (σ) on the probit scale (equation 2). Since σ is unaffected by *K*_i_, it is fully comparable between seed lots. Unfortunately, the largest comparative studies of seed longevities (Probert et al., 2009; Merritt et al., 2014; Satyanti et al., 2018) do not report initial viabilities or publish underlying data and thus, it is not possible to calculate σ (but see Lee et al., 2019, 2020). It is thus impossible to verify whether results of these studies also hold without the noise caused by the variation in initial viabilities. To improve comparability Hay et al. (Hay et al., 2022) called for publishing the theoretical initial viabilities and σ values alongside the *p*_50_ estimates, yet this has been done only by a few authors so far (e.g., Kochanek et al., 2009; Carta et al., 2018; Hay et al., 2019; White et al., 2023; Balasupramaniyam et al., 2025).

In summary, the *p*_50_ estimate is a suboptimal measure of seed longevity in storage, because it contains substantial noise caused by variation in the initial viabilities of the seed lots at the start of an aging experiment. Below we suggest how to reduce the noise in *p*_50_ estimates and present two alternative measures that are not affected by the noise caused by varying initial viability.

### Solution 1: Use seed lots with a limited range of initial viabilities

One way to reduce the unintended variation in *p*_50_ estimates is to restrict the seed accessions that are used for the experiment to a defined range of initial viabilities. This will reduce the number of usable accessions, but also the noise in the resulting *p*_50_ estimates. If set sufficiently high, it will also reduce the likelihood of obtaining negative estimates of *p*_50_, which are also difficult to handle in subsequent analyses. Since this solution makes the fewest changes to the established practice of using *p*_50_ estimates, it might be most acceptable for some researchers. In this context, the least noise will be achieved when using only accessions with viability below 95%. While this might be counter intuitive, we show that the unintended variation in *p*_50_ estimates is approximately five times larger when using seeds with very high initial viabilities (> 90%) compared to 75-85% (Figure 3). So far, comparative studies on seed longevity have restricted the initial viabilities from the bottom (>75, 85 or >90%) but have never set an upper limit (e.g., Probert et al., 2009; Merritt et al., 2014; Satyanti et al., 2018). While restricting the range of initial viabilities to a specific range reduces the unintended variation in *p*_50_ estimates, the estimates still contain some noise, just less than the *p*_50_ estimates currently in use. We thus consider this approach less optimal than the two solutions presented below.

**Figure 3:**
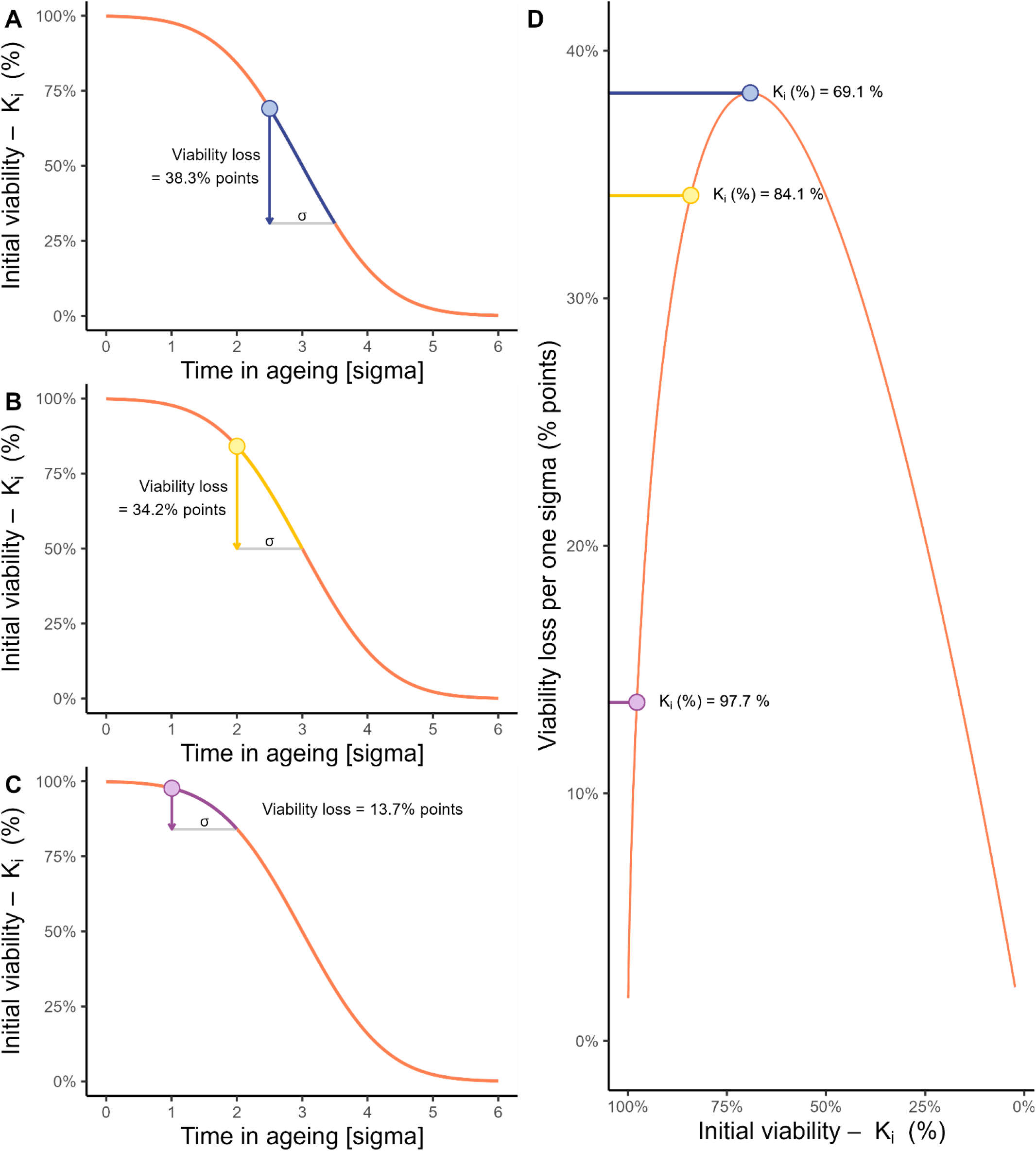
Illustration of loss of seed viability per one σ, that is the fixed number of days, and how this loss varies with the initial seed viability. A-C show percentage viability loss over one σ on the seed survival curve for three initial viabilities, D shows the magnitude of percentage viability loss per one σ as a function of initial viability.

### Solution 2: Use the rate of seed viability loss, σ

The probit rate of seed viability loss (σ) is the direct measure of seed longevity in storage and is independent of the initial viability. It represents the inverse of the slope of the probit generalized linear model used to describe the loss of seed viability in storage (Ellis and Roberts, 1980; Wolkis et al., 2025) and can be calculated using seeds with any reasonable initial viability. “Reasonable viability” in this context means a viability that enables reliable estimation of σ. If the initial viability is low and quickly drops to zero, there may be a lack of informative data points. Hay et al. (2022) already argued that σ is the optimal measure of seed longevity for comparative studies across species, accessions and genotypes, but acknowledged that it is difficult to interpret without understanding the probit equation. Nevertheless, σ was reported alongside *p*_50_ in some previous studies, but comparisons among accessions were mostly based on *p*_50_ (Hay et al., 2010; Moravcová et al., 2022, but see Hay and Probert, 1995; Kochanek et al., 2009; Carta et al., 2018).

The probit or “probability unit”, is rather abstract. In the context of seed aging, one would intuitively expect to translate σ to days in aging that are required for seeds to lose viability by a certain proportion. However, the rate of viability loss in percentage terms is not linear over time (Hay et al., 2019). Instead, the different viability intervals of for example 84.1 to 50%, 97.7 to 84.1%, and 99.9 to 99.7% all correspond to differences of one probit. In other words, a viability decline from 99.9 to 97.7%, or from 97.7 to 84.1%, or from 84.1 to 50% all take the same number of days, described by σ (Figure 3). The loss of percentage viability is steepest in the middle of the curve, and the largest drop in percentage viability for one probit unit is from 69 to 31%, i.e., from +0.5 to -0.5 on the probit scale. In simple terms, σ can be described as the time needed for viability to fall from e.g., 84 to 50% (Hay and Probert, 1995). The σ can be obtained directly from the seed viability equation, as the inverse of the slope of the probit curve (specifically, -1/slope). The calculation for σ from the survival curve is included in the annotated R-script in the supplementary materials (Supplementary Material 2)

### Solution 3: Adjust p_50_ to a standardized initial viability

A third option is to use the rate of seed viability loss (σ) to calculate *p*_50_ standardized to a certain theoretical initial viability, e.g., 90%. This approach allows the use of any seed accession for which it is possible to estimate σ, and then calculate how many days it would take for viability (of this accession) to fall from 90 to 50%. This calculation is very simple, and we provide an annotated R-script in the supplementary materials (Supplementary Material 2). This standardized *p*_50_ is proportional to σ, without any unintended noise. The standardized *p*_50_ could be more accessible as the concept is close to the *p*_50_ estimates in current practice while avoiding the unintended noise introduced by varying initial viabilities.

### Final remarks

Regardless of the parameter chosen to compare seed longevity in storage among accessions in a given study, it is essential to report all parameters of the seed viability curve - the theoretical initial viability (*K*_i_) and the probit rate of seed viability loss (σ). This will enable reciprocal recalculation among the estimates of *p*_50,_ σ and standardized *p*_50_ and allow comparisons in systematic reviews and meta-analyses. Our supplementary materials include an annotated R-script for these calculations (Supplementary Material 2).

All parameters should be accompanied by confidence intervals. While reporting uncertainties was not the focus of this study, we emphasize that real experimental data are always affected by some noise and this causes uncertainty in the estimated parameters. Standard errors of the parameters estimated from the model are symmetrical around the mean only on probit scale as provided in the model output, and become non-symmetrical when back-transformed to meaningful units. For example, a slope of -0.05 ± 0.01 S.E. corresponds to σ=20 days, with a confidence interval of 16.67 to 25 days, note that it is asymmetrical around the σ. Past studies on wild species mostly did not publish any uncertainties, only a few of them reported standard errors of the estimates, sometimes back-transformed (e.g., Hay and Probert, 1995; Carta et al., 2018; Hay et al., 2019; White et al., 2023; Balasupramaniyam et al., 2025). In the supplementary material (Supplementary Material 2), we provide an annotated R-code for the calculation of confidence intervals for all parameters.

Last but not least, we would like to encourage authors of future studies on seed longevity in storage to publish the raw germination data and experimental designs (Hay et al., 2022) and to share their data within open access initiatives (e.g., SeedArc, Fernández-Pascual et al., 2023). This will ensure that their data are preserved and re-used not only in the context of seed aging but also to explore the evolutionary and ecological drivers of seed germination.

## References

Balasupramaniyam, S., D. J. Merritt, F. R. Hay, and E. L. Dalziell. 2025. Assessing the storage potential of seed collections to inform the management of wild species seed banks L. Prior [ed.],. Australian Journal of Botany 73.

Carta, A., S. Bottega, and C. Spanò. 2018. Aerobic environment ensures viability and anti-oxidant capacity when seeds are wet with negative effect when moist: implications for persistence in the soil. Seed Science Research 28: 16–23.

Corbineau, F. 2024. The Effects of Storage Conditions on Seed Deterioration and Ageing: How to Improve Seed Longevity. Seeds 3: 56–75.

Ellis, R. H., and E. H. Roberts. 1980. Improved Equations for the Prediction of Seed Longevity. Annals of Botany 45: 13–30.

Fawcett, T. W., and A. D. Higginson. 2012. Heavy use of equations impedes communication among biologists. Proceedings of the National Academy of Sciences 109: 11735–11739.

Fernández-Pascual, E., A. Carta, S. Rosbakh, L. Guja, S. S. Phartyal, F. A. O. Silveira, S. C. Chen, et al. SeedArc, a global archive of primary seed germination data. New Phytologist 240: 466– 470.

Hay, F. R., R. M. Davies, J. B. Dickie, D. J. Merritt, and D. M. Wolkis. 2022. More on seed longevity phenotyping. Seed Science Research 32: 144–149.

Hay, F. R., A. Mead, and M. Bloomberg. 2014. Modelling seed germination in response to continuous variables: use and limitations of probit analysis and alternative approaches. Seed Science Research 24: 165–186.

Hay, F. R., D. J. Merritt, J. A. Soanes, and K. W. Dixon. 2010. Comparative longevity of Australian orchid (Orchidaceae) seeds under experimental and low temperature storage conditions. Botanical Journal of the Linnean Society 164: 26–41.

Hay, F. R., and R. J. Probert. 1995. Seed Maturity and the Effects of Different Drying Conditions on Desiccation Tolerance and Seed Longevity in Foxglove (Digitalis purpurea L.). Annals of Botany 76: 639–647.

Hay, F. R., R. Valdez, J.-S. Lee, and P. C. Sta. Cruz. 2019. Seed longevity phenotyping: recommendations on research methodology. Journal of Experimental Botany 70: 425–434.

Kochanek, J., K. J. Steadman, R. J. Probert, and S. W. Adkins. 2009. Variation in seed longevity among different populations, species and genera found in collections from wild Australian plants. Australian Journal of Botany 57: 123–131.

Lee, J.-S., J. Kwak, and F. R. Hay. 2020. Genetic markers associated with seed longevity and vitamin E in diverse Aus rice varieties. Seed Science Research 30: 133–141.

Lee, J.-S., M. Velasco-Punzalan, M. Pacleb, R. Valdez, T. Kretzschmar, K. L. McNally, A. M. Ismail, et al. 2019. Variation in seed longevity among diverse Indica rice varieties. Annals of Botany 124: 447–460.

Merritt, D. J., A. J. Martyn, P. Ainsley, R. E. Young, L. U. Seed, M. Thorpe, F. R. Hay, et al. 2014. A continental-scale study of seed lifespan in experimental storage examining seed, plant, and environmental traits associated with longevity. Biodiversity and Conservation 23: 1081–1104.

Mondoni, A., R. J. Probert, G. Rossi, E. Vegini, and F. R. Hay. 2011. Seeds of alpine plants are short lived: implications for long-term conservation. Annals of Botany 107: 171–179.

Moravcová, L., A. Carta, P. Pyšek, H. Skálová, and M. Gioria. 2022. Long-term seed burial reveals differences in the seed-banking strategies of naturalized and invasive alien herbs. Scientific Reports 12: 8859.

Nadarajan, J., C. Walters, H. W. Pritchard, D. Ballesteros, and L. Colville. 2023. Seed Longevity—The Evolution of Knowledge and a Conceptual Framework. Plants 12: 471.

Newton, R., F. Hay, and R. Probert. 2014. Protocol for comparative seed longevity testing.

Priestley, D. A., V. I. Cullinan, and J. Wolfe. 1985. Differences in seed longevity at the species level. Plant, Cell & Environment 8: 557–562.

Probert, R. J., M. I. Daws, and F. R. Hay. 2009. Ecological correlates of ex situ seed longevity: a comparative study on 195 species. Annals of Botany 104: 57–69.

Royal Botanic Gardens, Kew. 2022. Comparative Seed Longevity. https://brahmsonline.kew.org/Content/Projects/msbp/resources/Training/01-Comparative-longevity.pdf

Satyanti, A., A. B. Nicotra, T. Merkling, and L. K. Guja. 2018. Seed mass and elevation explain variation in seed longevity of Australian alpine species. Seed Science Research 28: 319–331.

Solberg, S.Ø., F. Yndgaard, C. Andreasen, R. von Bothmer, I. G. Loskutov, and Å. Asdal. 2020. Long-Term Storage and Longevity of Orthodox Seeds: A Systematic Review. Frontiers in Plant Science 11.

Walters, C., L. M. Wheeler, and J. M. Grotenhuis. 2005. Longevity of seeds stored in a genebank: species characteristics. Seed Science Research 15: 1–20.

White, F. J., F. R. Hay, T. Abeli, and A. Mondoni. 2023. Two decades of climate change alters seed longevity in an alpine herb: implications for ex situ seed conservation. Alpine Botany 133: 11– 20.

Wolkis, D., A. Carta, S. Rezaei, and F. R. Hay. 2025. Seed longevity: analysing post-storage germination data in R to fit the viability equation. Seed Science Research: 1–8.

