## Supplementary Material 1 for "Limitations of *p*_50_ as a measure of seed longevity and the way forward"

### Limitations of p50 as a measure of seed longevity and the way forward (Script to generate results and figures)

Klepka, Lea
Bucharova, Anna

2025-05-16

#### Content:

(0) Basic Code for Probit analysis (useful to understand further code)

(1) Relationship p50 ~ Ki (Figure 1A,B)

(2) Noise in p50 due to differences in initial viability (Figure 1B,D)

(3) Solutions to reduce/remove the noise in p50
   (3.1) Restrict range of initial viabilities (Figure 2)
   (3.2) Use sigma instead of p50 (Figure 3)
   (3.3) Standardized p50 (as illustrated in Figure 2A)

#### Load packages

```
library(drc)
library(MASS)
library(dplyr)
library(ggplot2)
library(cowplot)
library(arm)
```

#### Set consistent sizes for plots

```
point.size = 0.8
line.size = 0.75
lab.size = 12
text.size = 8
```

### (0) BASIC CODE FOR VISUALIZING SEED LONGEVITY MODEL RESULTS

#### 0.1 Probit transformation and backtransformation

```
qnorm(0.999) # function to transform percentage probability to probit scale
```

```
[1] 3.090232
```

```
pnorm(-2) # function tor back-transformation from probit to percentage probability
```

```
[1] 0.02275013
```

```
# (pnorm(0)= 0.5)) 50% probability coresponds to a probit-value of 0
```

Note: A viability of 100% corresponds to "Inf" on the probit scale and must be avoided in computations.

#### 0.2 Relationship: Percentage - Probit

```
seed_viability <- seq(from=0.001, to=0.999, by=0.001) # generate vector with
                                                       # seed viability between 0.001 and 0.999

# generate dataframe with seed viability and corresponding probit value
Transformation <- data.frame(
  seed_viability = seed_viability,
  probit = qnorm(seed_viability)) # see qnorm() function above

# plot relationship between seed viability (%) and corresponding probit value
p_transformation <- ggplot(data = Transformation, aes(x = seed_viability, y = probit)) +
  geom_line(size = line.size, colour = "#FF7F50") +
  labs(x = "Seed viability", y = "Probit") +
  scale_x_continuous(labels = scales::percent) +
  scale_y_continuous(breaks = seq(-3,3,1)) +
  theme_classic()

p_transformation # show plot
```

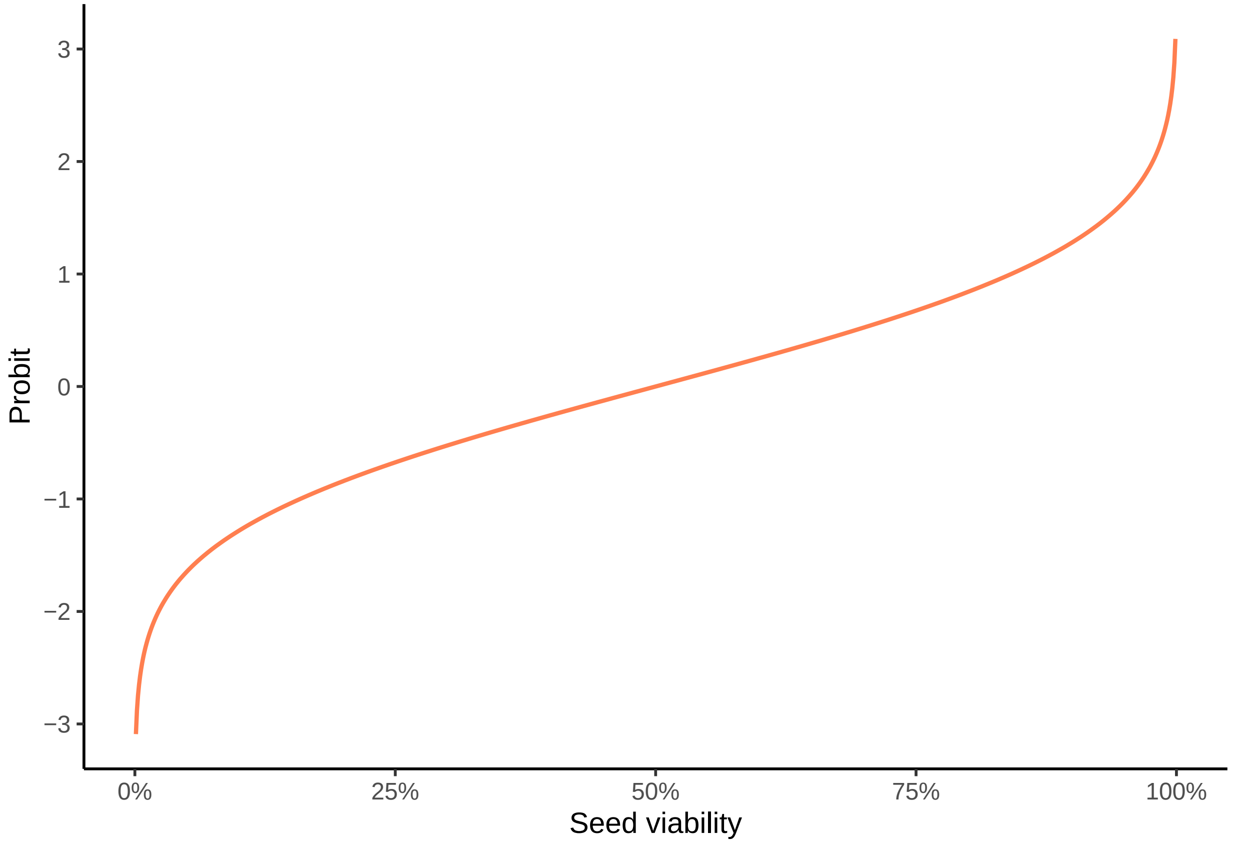


##### 0.3 Illustrate seed viability in storage in percentage and probits

###### Draw a curve that illustrates seed viability (%) in storage

Sigma = time in storage needed to reduce viability by 1 probit

```
ageing_in_sigma <- seq(from=3, to=-3, by=-0.01) # Percentage of germination success
                                                # range from 0.1349898% to 99.86501%
                                                # corresponds to probit range -3 to 3
time_in_sigma <- (-1*ageing_in_sigma)+3 # shift range from -3 to 3 (contains negative values)
                                        # to 0 to 6 (contains only positive values)
seed_viability <- pnorm(ageing_in_sigma) # calculate seed viability (back transformation from probit)
Ki <- qnorm(seed_viability) # probit transformation (from percentage to probit)

# generate dataframe to illustrate seed survival
Seed_survival <- data.frame(
  time_in_sigma = time_in_sigma,
  seed_viability = seed_viability)

# Plot Seed survival curve (relationship between seed viability (%) and time in ageing (sigma))
p_seed_survival <- ggplot(data = Seed_survival, aes(x = time_in_sigma, y = seed_viability)) +
  geom_line(size = line.size, colour = "#FF7F50") +
  labs(x = "Time in ageing [sigma]", y = expression("Seed viability (%)")) +
  scale_y_continuous(labels = scales::percent) +
  scale_x_continuous(breaks = seq(min(Seed_survival$time_in_sigma),
```

```
                                max(Seed_survival$time_in_sigma), 1)) +
  theme_classic() +
  theme(
    axis.title.x = element_text(size = lab.size),
    axis.title.y = element_text(size = lab.size),
    axis.text.x = element_text(size = text.size),
    axis.text.y = element_text(size = text.size),
    plot.margin = margin(c(0,0,0,0))
  )

p_seed_survival # show plot
```

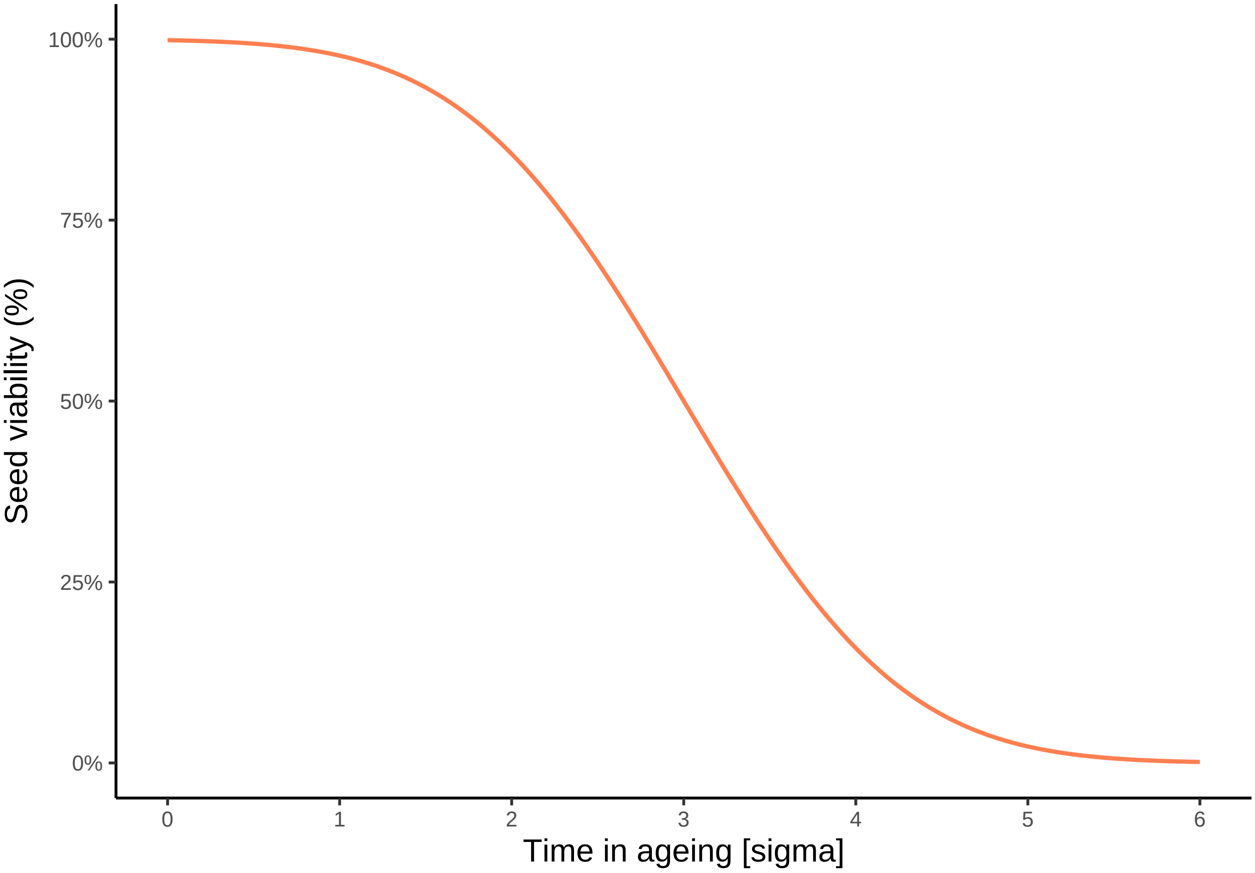


##### Draw a curve that illustrates seed viability (probits) in storage

Relationship between seed viability (probit) and time in ageing (sigma))

```
p_seed_survival_probit <- ggplot(data = Seed_survival,
                                aes(x = time_in_sigma, y = qnorm(seed_viability))) +
  geom_line(size = line.size, colour = "#FF7F50") +
  labs(x = "Time in ageing (sigma)", y = "Seed viability (probit)") +
  scale_x_continuous(breaks = seq(min(Seed_survival$time_in_sigma),
                                max(Seed_survival$time_in_sigma), 1)) +
  theme_classic() +
  theme(
```

```
    axis.title.x = element_text(size = lab.size),
    axis.title.y = element_text(size = lab.size),
    axis.text.x = element_text(size = text.size),
    axis.text.y = element_text(size = text.size),
    plot.margin = margin(c(0,0,0,0))
  )

p_seed_survival_probit # show plot
```

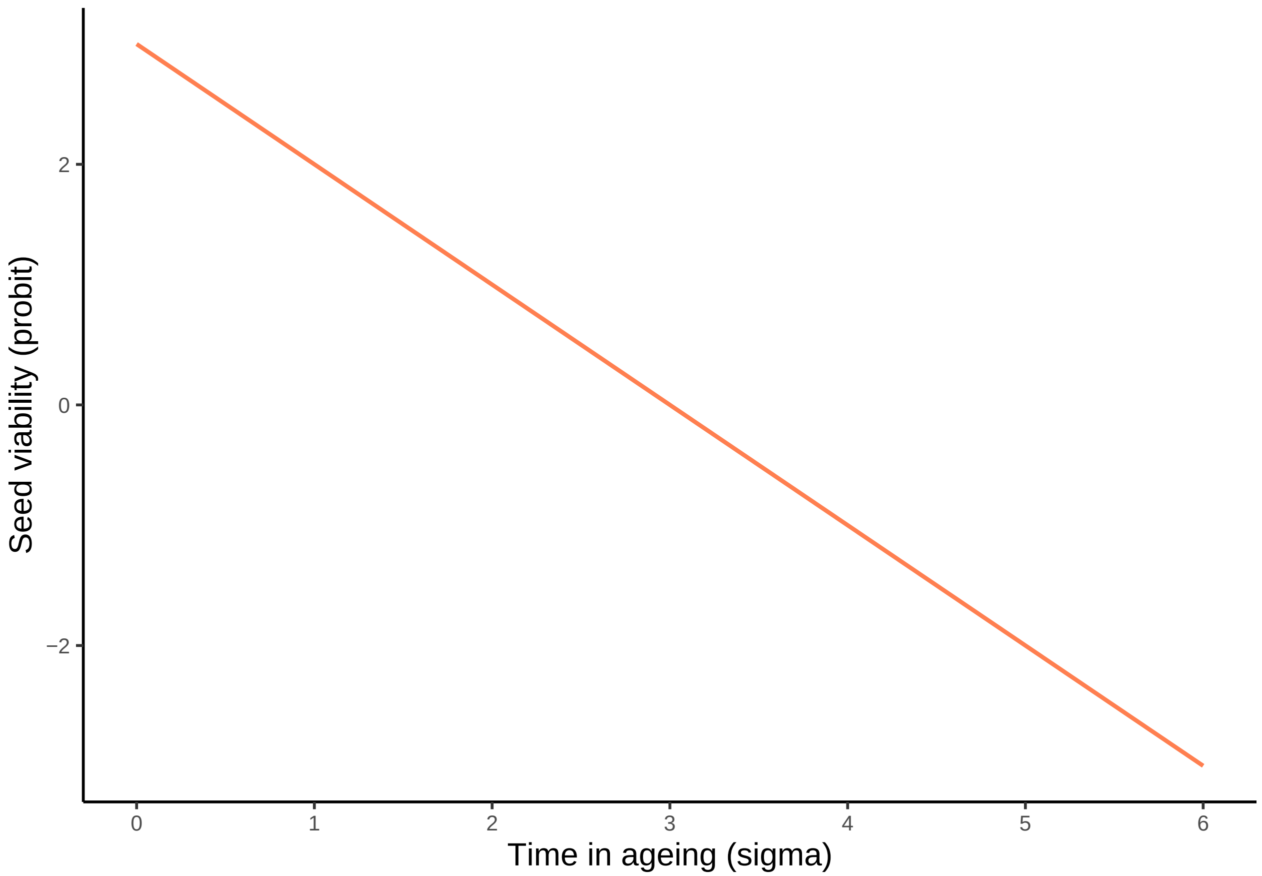


##### Combine seed survival plots to compare viability in percentage and probit

```
p_0 <- plot_grid(p_seed_survival, p_seed_survival_probit, ncol = 2,
                 labels = c("A", "B"), label_size = 12, label_x = 0, label_y = 1,
                 rel_widths = c(0.5, 0.5))

p_0 # show plot
```

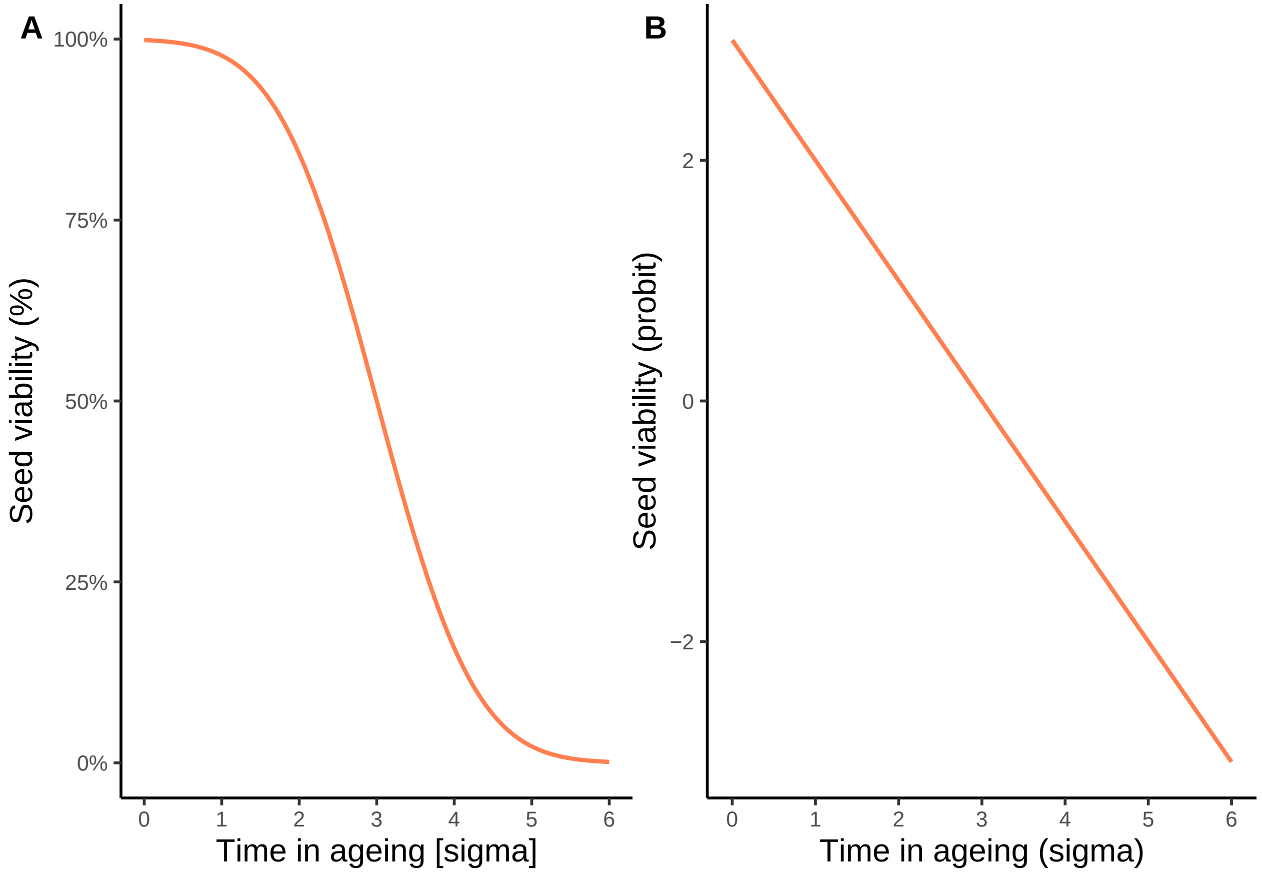


### (1) RELATIONSHIP BETWEEN P50 AND INITIAL VIABILITY (Ki)

#### 1.1 Simulate hypothetical species

```
sigma_days <- 20 # set fixed sigma in days
sigma_probit = -1/sigma_days # slope in probit units per day
                              # (after 1 day, probit decreases by 1/20)

viabilities <- seq(from=0.85, to=0.999, by=0.001) # sequence of initial viabilities (in percentage)
                                                   # from 0.85 to 0.999
Ki.probit <- qnorm(viabilities) # transformation of viabilities from percentage to probit
p50 <- Ki.probit*sigma_days # calculate P50 applying equation (2): p50 = Ki*sigma

# create dataframe for p50~Ki relationship
Relationship_p50_Ki <- data.frame(
  viability = viabilities,
  p50 = p50)
```

#### 1.2 Calculate minimum and maximum p50 and ratio

```
p50_min <- round(min(p50),2)
p50_max <- round(max(p50),2)
p50_ratio <- p50_max/p50_min

# show results
p50_min
```

```
[1] 20.73
```

```
p50_max
```

```
[1] 61.8
```

```
p50_ratio
```

```
[1] 2.981187
```

#### 1.3 Figure 1 B: Relationship between p50 and the initial seed viability

```
p_1B <- ggplot(data = Relationship_p50_Ki, aes(x = viability, y = p50)) +
  geom_line(colour = "#FF7F50", size=line.size) +
  labs(x= expression("Initial viability - "~ K[i]~" (%)"),
       y = expression(p[50] ~ " [days]")) +
  scale_x_continuous(labels = scales::percent, limits = c(0.85,1.01)) +
  scale_y_continuous(limits = c(0,70)) +
  theme_classic() +
  theme(
    axis.title.x = element_text(size = lab.size),
    axis.title.y = element_text(size = lab.size),
    axis.text.x = element_text(size = text.size),
    axis.text.y = element_text(size = text.size),
    plot.margin = margin(c(0,0,0,0))
  )

p_1B # show plot
```

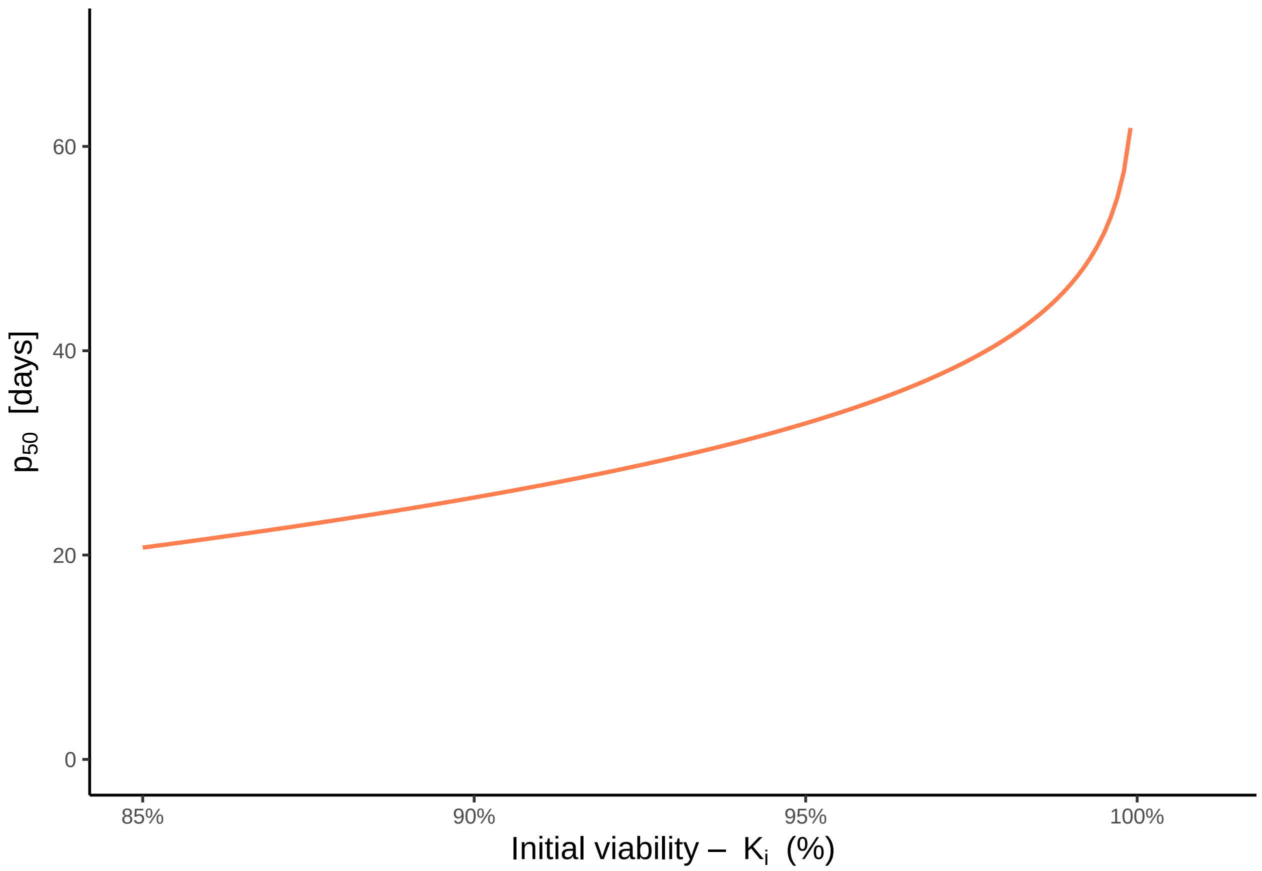


##### 1.4 Figure 1A: Seed viability loss

with sigma = 20 and 4 different initial viabilities. This code is rather long because it contains many design elements.

```
sigma_days <- 20 # set fixed sigma in days
sigma_probit = -1/sigma_days # slope in probit units per day (after 1 day, probit decreases by 1/20)

# generate dataframe for hypothetical seed ageing experiment
days_in_ageing <- seq(0,100,1) # sequence of ageing duration (between 0 and 100 days)
viability_probit <- qnorm(0.9991) + sigma_probit*days_in_ageing # calculate seed viability for each
                                                                # ageing duration in probit;
                                                                # Sigma_probit = viability loss per day
viability <- pnorm(viability_probit) # backtransformation to percentage

One_spec <- data.frame(
  days_in_ageing = days_in_ageing,
  viability = viability)

# generate an example p50 for 4 defined initial viabilities (Ki)
Ki_percentage <- c(0.85,0.90,0.95,0.999) # vector of initial viabilities (in percentage)
Ki_probit <- qnorm(Ki_percentage) # transformation of viabilities from percentage to probit
p50 <- Ki_probit*sigma_days # calculate P50 applying equation (2):
# p50 = Ki*sigma (one p50 value for each initial viability)
```

```
curve_start <- 0.9991 # theoretical maximum viability
# --> this is where the seed survival curve starts (the same for all initial viabilities)
# calculate the number of days it takes to decay from theoretical maximum viability (curve_start)
# to given initial viability - for plotting
x_viabs <- (qnorm(curve_start)-qnorm(Ki_percentage))*sigma_days # seed viability equation applied:
                                                                # p = (Ki-viability)*sigma

# define ends of grey lines for plotting
y_ends <- c(0.5, 0.43, 0.36, 0.29)

# labels - how long it takes to decay to p50 from given initial viability
p50lab <- paste0("p[50] == ", round(qnorm(Ki_percentage)*sigma_days,1), " * ' days'")
# calculation of p50 for given initial viability
viabslab <- paste0("K[i]~'(%)' == ", Ki_percentage*100, " * '%'")

# plot relationship between seed viability an ageing duration
p_1A <- ggplot(data = One_spec, aes(x=days_in_ageing, y=viability)) +
  geom_point(shape = 16, y = 0.5, x = qnorm(curve_start)*sigma_days,
             col="#E66A3A", size = 4) + # point to 50% germination (p50 = Ki*Sigma)
  geom_segment(aes(x=qnorm(curve_start) * sigma_days, xend=qnorm(curve_start) * sigma_days,
                   y=0.5,yend=min(y_ends)), color="grey80") + # vertical line at p50
  geom_line(size=line.size) # seed survival curve

# add labels and lines for each of the example initial viabilities
for (i in 1:length(Ki_percentage)) {
  segment_data <- data.frame(x = x_viabs[i], # how long it takes to reach given initial viability
                             y = y_ends[i], # lower ends of vertical lines
                             yend = Ki_percentage[i]-0.002, # upper end of vertical lines
                             p50 = qnorm(curve_start) * sigma_days, # p50 = Ki* sigma
                             labs = p50lab[i], # labels
                                               # (how long it takes to decay to p50 from
                                               # given initial viability)
                             xtext = 51.5, # Position of text
                             ytext = y_ends[i] + 0.02, # Position of text (0.03 above arrow line)
                             labsK = viabslab[i], # label for Ki(%)
                             xK = x_viabs[i], # Position of given initial viability
                             yK = Ki_percentage[i] + 0.02 ) # Position of given initial viability
                                                            # (at y=initial viability -0.08 down)
  p_1A <- p_1A +
    geom_segment(data = segment_data, aes(x = x, xend = x,
                                          y = y, yend = yend),
                 color = "grey80", size = line.size) + # vertical lines for
                                                       # each given initial viability
    geom_segment(data = segment_data, aes(x = x, xend = p50,
                                          y = y, yend = y),
                 color="#828282", size = 0.6,
                 arrow = arrow(ends = "both",length=unit(0.02,"npc"))) + # horizontal lines + arrow
    geom_text (data = segment_data, aes(x = xtext, y = ytext, label = labs),
               parse = T, size = text.size, size.unit = "pt") + # text: p50 = 20.7 days, ...
    geom_text (data = segment_data, aes(x = xK, y = yK, label=labsK),
               hjust = 0, parse = T, size = text.size, size.unit = "pt") # text: given initial
                                                                          # viability
}
```

```r
p_1A <- p_1A +
  labs(x = "Days in ageing", y = expression("Initial viability - "~ K[i]~" (%)")) +
  scale_y_continuous (labels = scales::percent) +
  theme_classic() +
  theme(
    axis.title.x = element_text(size = lab.size),
    axis.title.y = element_text(size = lab.size),
    axis.text.x = element_text(size = text.size),
    axis.text.y = element_text(size = text.size),
    plot.margin = margin(c(0,0,0,0))
  )

p_1A # show plot
```

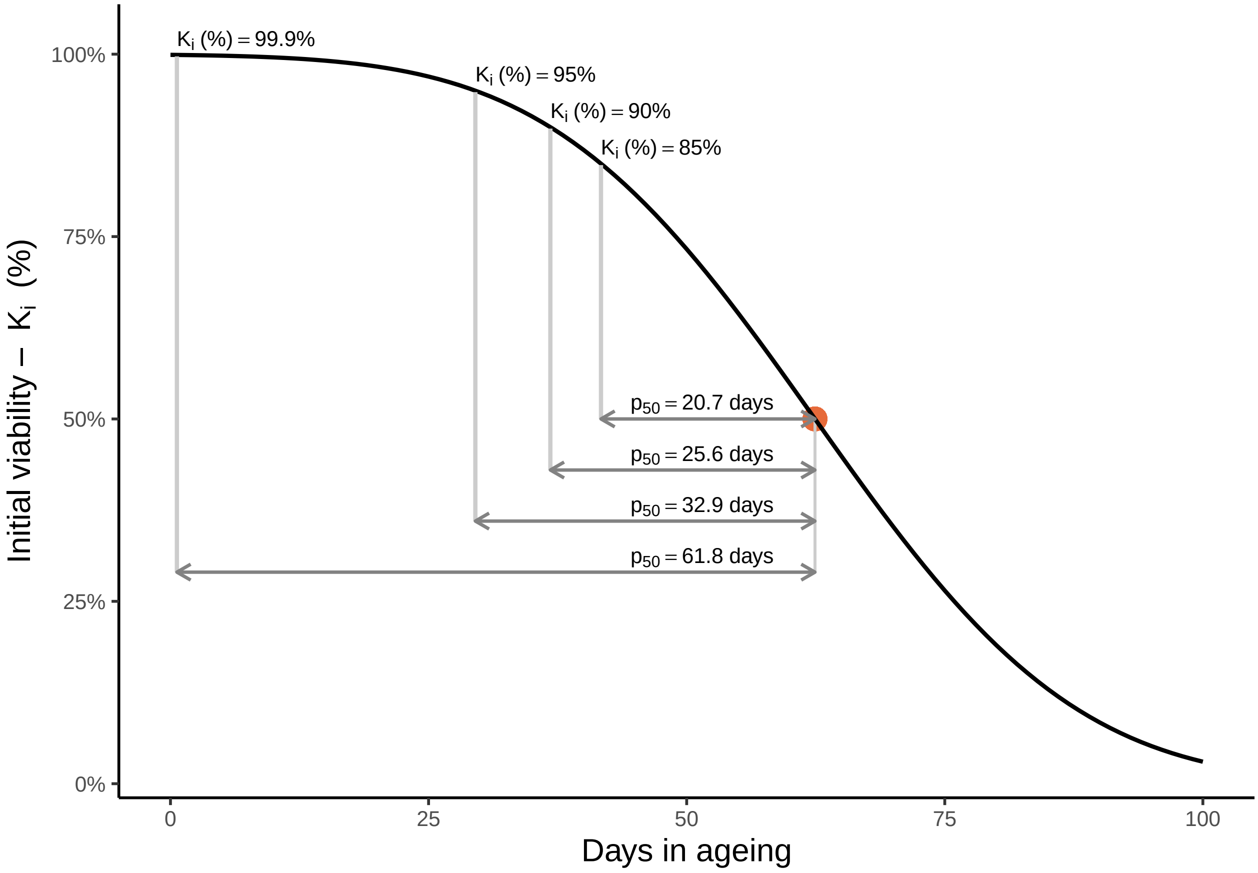


### (2) NOISE IN P50 DUE TO DIFFERENCES IN INITIAL VIABILITY

#### 2.1 Simulate 500 seed lots with random initial viabilities and random sigma-values

```
lot_number <- 500 # number of random seed lots
viability <- runif(lot_number, min = 0.85, max = 0.999) # random initial viabilities for
                                                        # each seed lot varying between
                                                        # min (0.85) and max (0.999)
sigma <- runif(lot_number, min = 1, max = 50) # random values of sigma (in days) for
                                              # each seed lot varying between
                                              # min (1 day) and max (50 days)

# create dataframe for simulated 500 random seed lots with generated initial viabilities and sigma
Simdat <- data.frame(
  viability = viability,
  sigma = sigma)
Simdat$ID <- row(Simdat)[,1] # assign IDs to simulated species

# calculate p50 for simulated seed lots with original initial viability and fixed Ki
Simdat <- Simdat %>%
  mutate(
    p50 = qnorm(viability) * sigma,
    p50_0.75 = qnorm(0.75) * sigma,  # Fixed Ki(%) at 0.75
    p50_0.80 = qnorm(0.80) * sigma,  # Fixed Ki(%) at 0.80
    p50_0.85 = qnorm(0.85) * sigma,  # Fixed Ki(%) at 0.85
    p50_0.90 = qnorm(0.90) * sigma,  # Fixed Ki(%) at 0.90
    p50_0.95 = qnorm(0.95) * sigma,  # Fixed Ki(%) at 0.95
    p50_0.999 = qnorm(0.999) * sigma # Fixed Ki(%) at 0.999
  )
```

#### Colour gradients for plots

```
# function for plots
create_gradient <- function(low, high) {
  scale_color_gradient2(
    low = "#3B4994", mid = "#FF7F50", high = "#FFC300",
    midpoint = 0.93, limits = c(low, high)
  )
}

# function for legend in Figure 1
create_gradient_label <- function(low, high) {
  scale_fill_gradient2(
    low = "#3B4994", mid = "#FF7F50", high = "#FFC300",
    midpoint = 0.93, limits = c(low, high)
  )
}

# create dummy dataframe for gradient legend (Figure 1)
Dummy_data <- data.frame(
  x = rep(seq(min(Simdat$viability), max(Simdat$viability),
              length.out = 100), each = 100),  # Create a grid for x
  y = rep(seq(min(Simdat$viability), max(Simdat$viability),
              length.out = 100), times = 100)  # Create a grid for y
)
```

```r
# Create legend (Figure 1)
p_legend_gradient <- ggplot(Dummy_data, aes(x = x, y = y, fill = y)) +
  geom_tile() +  # create a grid of rectangles
  create_gradient_label(min(Simdat$viability), max(Simdat$viability)) + # create gradient
  ylim(min(Simdat$viability), 1.2) +
  scale_y_continuous(labels = scales::percent, name = NULL, position = "right") + # Remove y-axis label
  scale_x_continuous(
    name = expression("Initial viability - " ~ K[i] ~ "(%)") ,breaks = NULL, # Remove x-axis labels
                                                                        # and ticks
    position = "top") +
  theme_classic() +
  theme(
    axis.text.y = element_text(size = text.size),  # Hide axis text
    axis.text.x = element_blank(),  # Hide axis text
    axis.title.x = element_text(size = text.size, hjust = 0),
    axis.ticks = element_line(),  # Keep the ticks but remove the axis line
    axis.ticks.y = element_line(),  # Keep y-axis ticks
    axis.line = element_blank(),  # Remove the axis line
    panel.border = element_blank(),  # Remove the panel border
    legend.position = "none",
    plot.margin = margin(80,5,80,10),
  )

p_legend_gradient # show legend
```

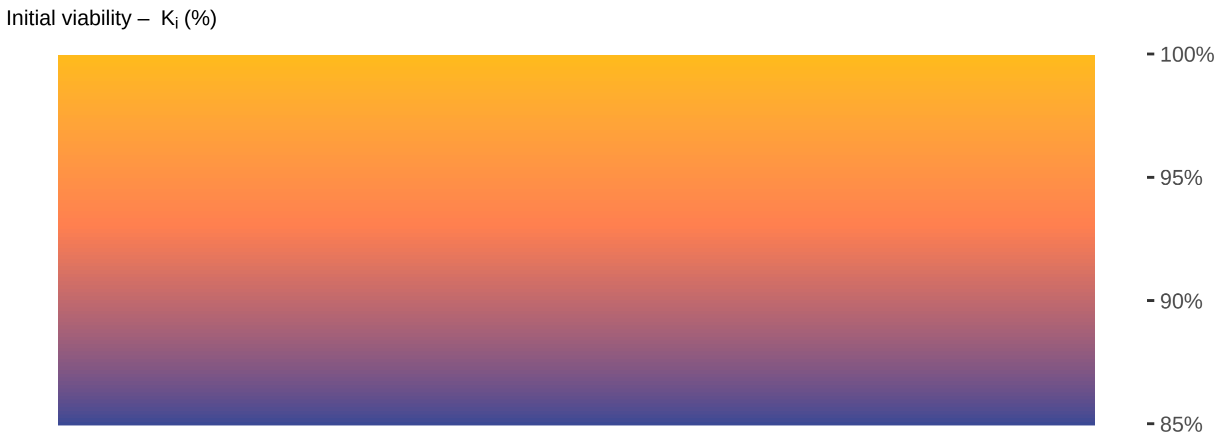

#### 2.2 Figure 1C: Noise in p50

using seed lots with initial viabilities varying between 85 and 99.9%

```
p_1C <- ggplot(data = Simdat, aes(x = sigma, y = p50, color = viability)) +
  geom_line(aes(x = sigma, y = p50_0.85), color = "#828282", size = line.size) +
  geom_line(aes(x = sigma, y = p50_0.999), color = "#828282", size = line.size) +
  geom_point(shape = 19, alpha = 0.6, size = point.size) +
  create_gradient(min(Simdat$viability), max(Simdat$viability)) +  # Use whole gradient
  ylim(0, 150) +
  labs(x = "Sigma", y = expression(p[50] * " [days]")) +
  theme_classic() +
  theme(
    axis.title.x = element_text(size = lab.size),
    axis.title.y = element_text(size = lab.size),
    axis.text.x = element_text(size = text.size),
    axis.text.y = element_text(size = text.size),
    plot.margin = margin(c(0,0,0,0))
  )

p_1C # show plot
```

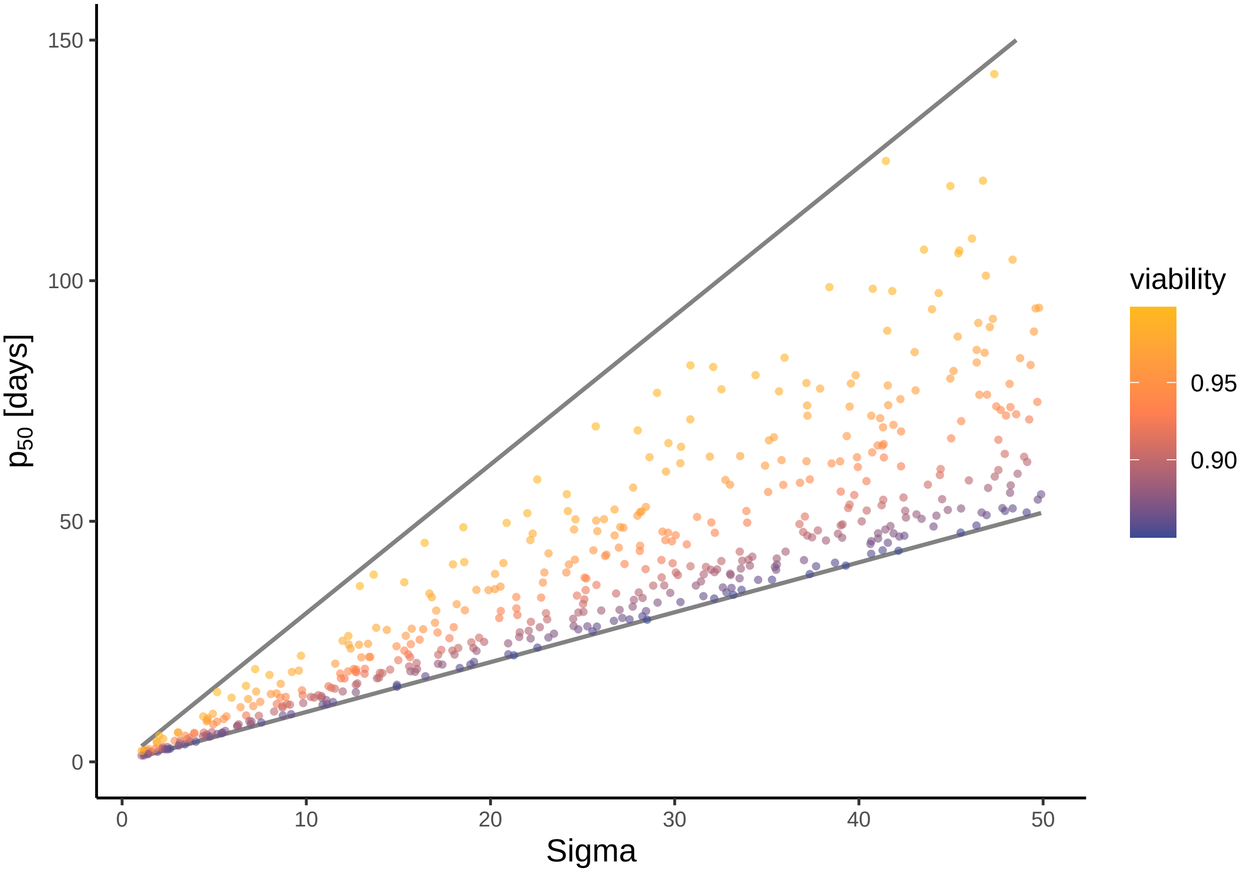


#### 2.3 Figure 1D: Log-transformation of p50

Same as Figure 1C, but with log-transformed y-axis

```r
p_1D <- ggplot(data = Simdat, aes(x = sigma, y = p50, color = viability)) +
  geom_line(aes(x = sigma, y = p50_0.85), color = "#828282", size = line.size) +
  geom_line(aes(x = sigma, y = p50_0.999), color = "#828282", size = line.size) +
  geom_point(shape = 19, alpha = 0.6, size = point.size) +
  create_gradient(min(Simdat$viability), max(Simdat$viability)) +  # Use whole gradient
  ylim(0, 150) +
  scale_y_log10(breaks = c(0, 0.1, 10, 100, 150)) +
  labs(x = "Sigma", y = expression(log(p[50]) * " [days]")) +
  theme_classic() +
  theme(
    axis.title.x = element_text(size = lab.size),
    axis.title.y = element_text(size = lab.size),
    axis.text.x = element_text(size = text.size),
    axis.text.y = element_text(size = text.size),
    plot.margin = margin(c(0,0,0,0))
  )

p_1D # show plot
```

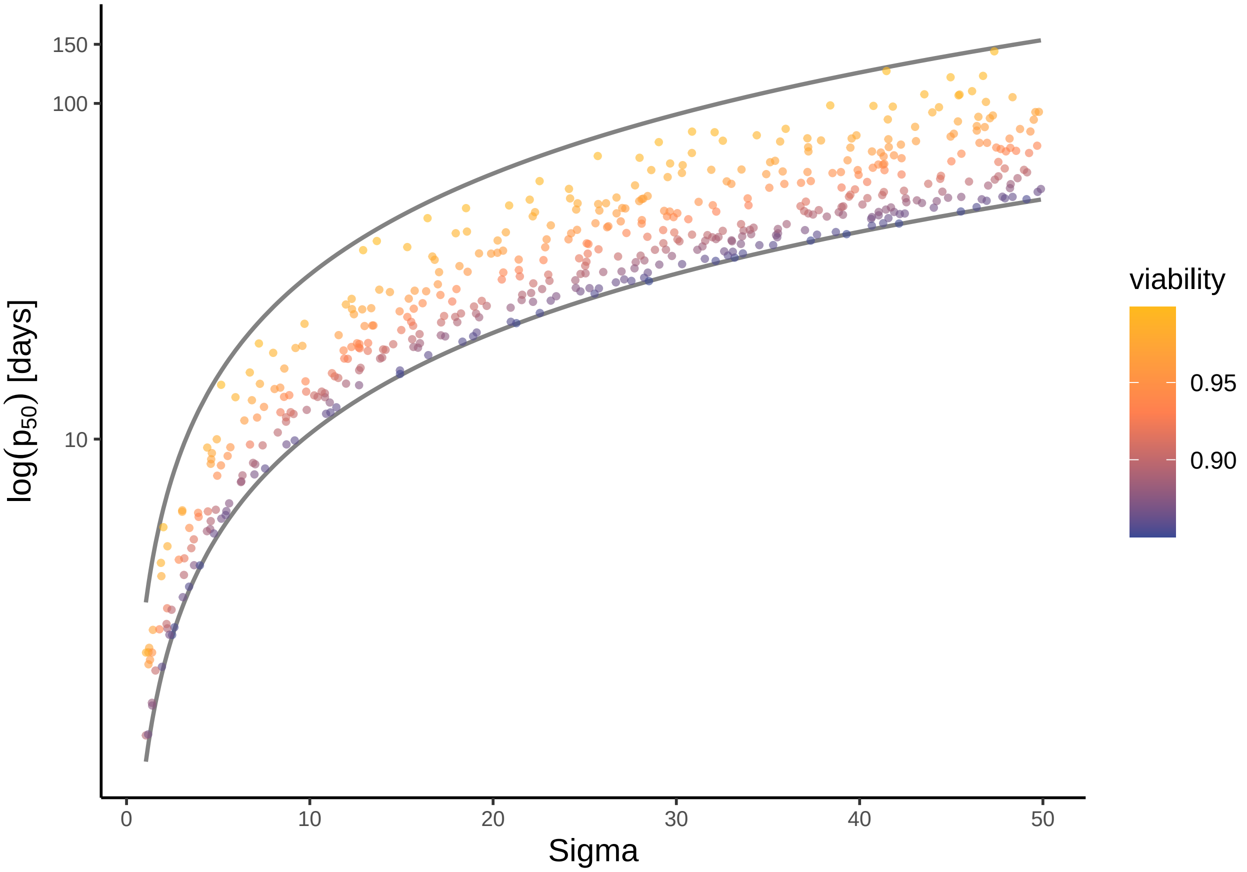


```r
# the relationship between p50 (if Ki(%)=0.85) and p50 (if Ki(%)=0.999)
unique(Simdat$p50_0.999/Simdat$p50_0.85)
```

```
[1] 2.981602 2.981602 2.981602
```

```r
# the relationship between p50 (if Ki=0.85) and p50 (if Ki=0.999), on log-scale
unique(log(Simdat$p50_0.999)-log(Simdat$p50_0.85))
```

```
[1] 1.092461 1.092461 1.092461 1.092461 1.092461
```

#### 2.4 Combine Figure 1

```r
# Remove legend from p_2C and p_2D
p_1C_nolegend <- p_1C + theme(legend.position = "none")
p_1D_nolegend <- p_1D + theme(legend.position = "none")

p_1 <- plot_grid(
  plot_grid(p_1A, p_1B, ncol = 2,
            labels = c("A", "B"), label_size = 12, label_x = 0, label_y = 1,
            rel_widths = c(0.7, 0.3)),  # Adjust the width ratio for the first row
  plot_grid(p_1C_nolegend, p_1D_nolegend, p_legend_gradient, ncol = 3,
            labels = c("C", "D", ""), label_size = 12, label_x = 0, label_y = 1.01,
            rel_widths = c(0.42, 0.42, 0.16)),
  nrow = 2
) + theme(plot.margin = margin(0,15,0,0))

p_1 # show plot
```

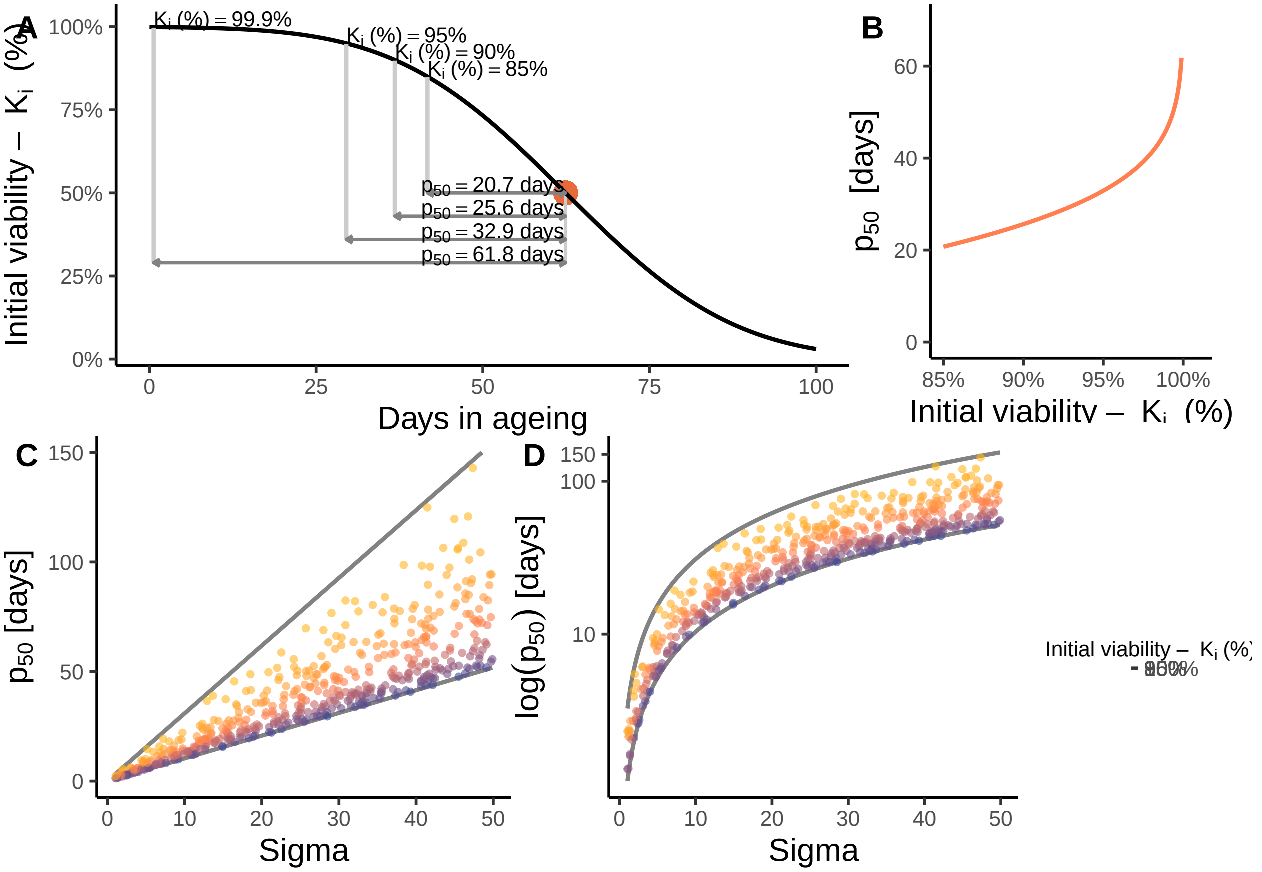

```
# save plot (the saved plots look identical to the figures in the paper)
# ggsave(file = "your/path/Figure1.pdf", plot = p_1, width = 180, height = 200, units = "mm", dpi=500) #
```

### (3) SOLUTIONS TO REDUCE/REMOVE THE NOISE IN P50

#### 3.1 Figure 2: Restrict range of initial viabilities

```
# simulate 500 seed lots with random initial viabilities and random sigma-values (same as in (2))
lot_number <- 500 # number of random seed lots
sigma <- runif(lot_number, min = 1, max = 50) # random values of sigma (in days) for each seed lot
                                              # varying between min (1 day) and max (50 days)

# create dataframe for simulated 500 random seed lots with generated initial viabilities and sigma
Simdat <- data.frame(sigma = sigma)
Simdat$ID <- row(Simdat)[,1] # assign IDs to simulated species

# calculate p50 for simulated seed lots that they would have had if Ki(%) was fixed
Simdat <- Simdat %>%
  mutate(
    p50_0.75 = qnorm(0.75) * sigma,  # Fixed Ki(%) at 0.75
    p50_0.80 = qnorm(0.80) * sigma,  # Fixed Ki(%) at 0.80
    p50_0.85 = qnorm(0.85) * sigma,  # Fixed Ki(%) at 0.85
    p50_0.90 = qnorm(0.90) * sigma,  # Fixed Ki(%) at 0.90
    p50_0.95 = qnorm(0.95) * sigma,  # Fixed Ki(%) at 0.95
    p50_0.999 = qnorm(0.999) * sigma # Fixed Ki(%) at 0.999
  )
```

##### Illustrate the relationship between sigma and p50 if initial viability is fixed

```
p_2A.1 <- ggplot(data = Simdat, aes(x = sigma, y = p50)) +
  geom_line(aes(x = sigma, y = p50_0.75), color = "#828282", size = line.size) + # for Ki(%) = 0.75
  geom_line(aes(x = sigma, y = p50_0.80), color = "#828282", size = line.size) + # for Ki(%) = 0.80
  geom_line(aes(x = sigma, y = p50_0.85), color = "#828282", size = line.size) + # for Ki(%) = 0.85
  geom_line(aes(x = sigma, y = p50_0.90), color = "#828282", size = line.size) + # for Ki(%) = 0.90
  geom_line(aes(x = sigma, y = p50_0.95), color = "#828282", size = line.size) + # for Ki(%) = 0.95
  geom_line(aes(x = sigma, y = p50_0.999), color = "#828282", size = line.size) + # for Ki(%) = 0.999

  annotate("text", x = 52, hjust = 0, y = max(Simdat$p50_0.75),
           label = expression(K[i] ~ "(%)" == 75 * "%"),
           size = text.size, size.unit = "pt") + # text for Ki(%) = 0.75
  annotate("text", x = 52, hjust = 0, y = max(Simdat$p50_0.80),
           label = expression(K[i] ~ "(%)" == 80 * "%"),
           size = text.size, size.unit = "pt") + # text for Ki(%) = 0.80
  annotate("text", x = 52, hjust = 0, y = max(Simdat$p50_0.85),
           label = expression(K[i] ~ "(%)" == 85 * "%"),
           size = text.size, size.unit = "pt") + # text for Ki(%) = 0.85
  annotate("text", x = 52, hjust = 0, y = max(Simdat$p50_0.90),
           label = expression(K[i] ~ "(%)" == 90 * "%"),
```

```
           size = text.size, size.unit = "pt") + # text for Ki(%) = 0.90
  annotate("text", x = 52, hjust = 0, y = max(Simdat$p50_0.95),
           label = expression(K[i] ~ "(%)" == 95 * "%"),
           size = text.size, size.unit = "pt") + # text for Ki(%) = 0.95
  annotate("text", x = 52, hjust = 0, y = max(Simdat$p50_0.999),
           label = expression(K[i] ~ "(%)" == 99.9 * "%"),
           size = text.size, size.unit = "pt") + # text for Ki(%) = 0.999
  xlim(0, 65) +
  labs(x = "Sigma", y = expression(p[50])) +
  theme_classic() +
  theme(
    axis.title.x = element_text(size = lab.size),
    axis.title.y = element_text(size = lab.size),
    axis.text.x = element_text(size = text.size),
    axis.text.y = element_text(size = text.size),
    plot.margin = margin(c(0,0,0,0))
  )

p_2A.1 # show plot
```

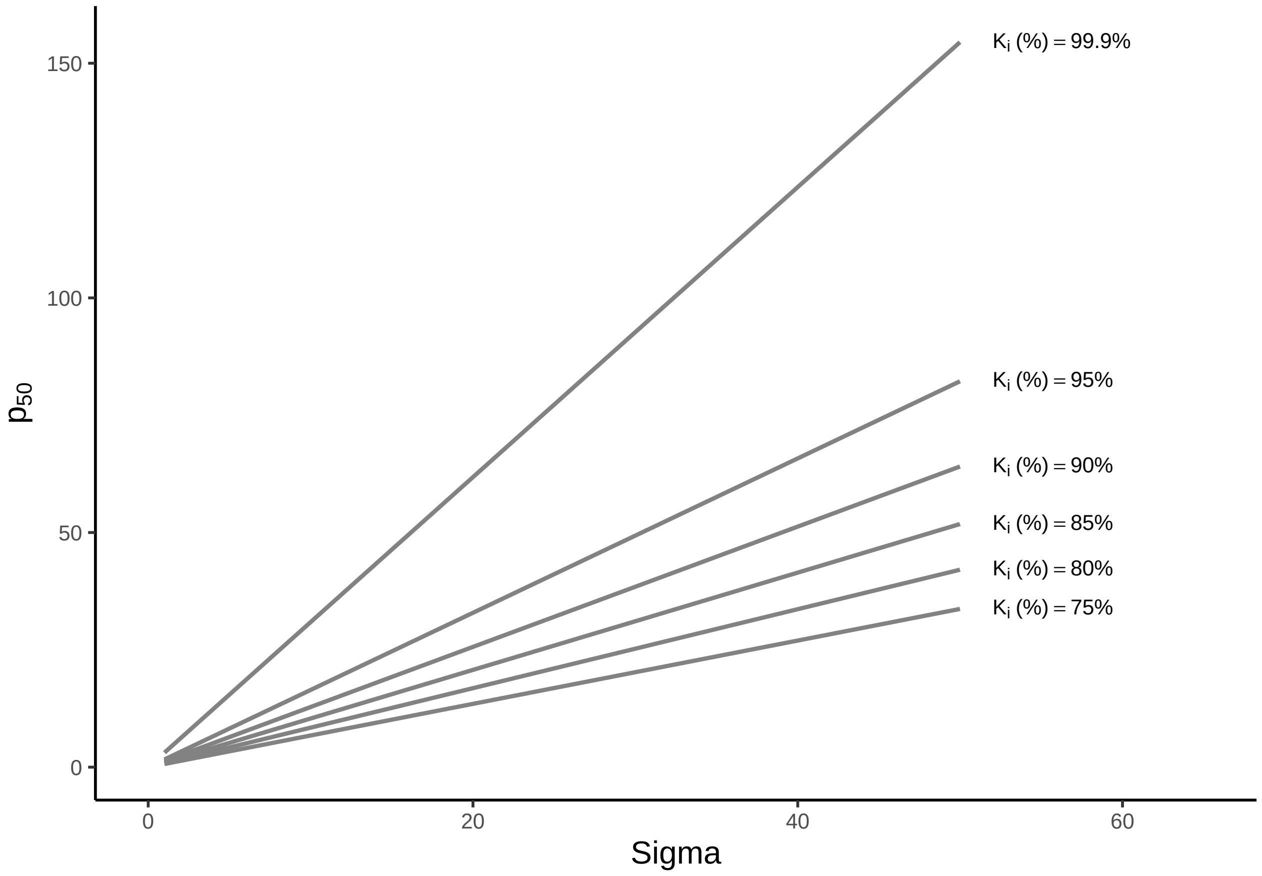

##### Illustrate the relationship between sigma and p50 if initial viability is limited to intervals

```
# interval Ki(%) 0.75-0.85
Ki_0.85_0.75 <- round(((Simdat$p50_0.85-Simdat$p50_0.75)/Simdat$sigma)[1], 2)
# interval Ki(%) 0.80-0.90
Ki_0.90_0.80 <- round(((Simdat$p50_0.90-Simdat$p50_0.80)/Simdat$sigma)[1], 2)
# interval Ki(%) 0.85-0.95
Ki_0.95_0.85 <- round(((Simdat$p50_0.95-Simdat$p50_0.85)/Simdat$sigma)[1], 2)
# interval Ki(%) 0.90-0.999
Ki_0.999_0.90 <- round(((Simdat$p50_0.999-Simdat$p50_0.90)/Simdat$sigma)[1], 2)

# generate interval information for Figure 2 (text on the right in Figure 1A)
p_2A.2 <- ggplot() +
  annotate("text", x = 1, y = 5.1,
           label = expression(bold("Intervals of "*K[i])),
           size = text.size, hjust = 0, size.unit = "pt") +
  annotate("text", x = 1, y = 4.9,
           label = expression(bold("(% and probit)")),
           size = text.size, hjust = 0, size.unit = "pt") +

  annotate("text", x = 2, y = 5.1, label =  expression(bold("Variability in " * p[50])),
           size = text.size, hjust = 0, size.unit = "pt") +
  annotate("text", x = 2, y = 4.9, label =  expression(bold("(fraction of " * sigma * ")")),
           size = text.size, hjust = 0, size.unit = "pt") +

  annotate("text", x = 1, y = 4.1, label = expression("75 - 85%"),
           size = text.size, hjust = 0, size.unit = "pt") +
  annotate("text", x = 1, y = 3.9, label = paste(round(qnorm(0.75),2),"-", round(qnorm(0.85),2)),
           size = text.size, hjust = 0, size.unit = "pt") +
  annotate("text", x = 2, y = 4, label = paste(Ki_0.85_0.75),
           size = text.size, hjust = 0, size.unit = "pt") +

  annotate("text", x = 1, y = 3.6, label = "80 - 90%",
           size = text.size, hjust = 0, size.unit = "pt") +
  annotate("text", x = 1, y = 3.4, label = paste(round(qnorm(0.8),2),"-", round(qnorm(0.9),2)),
           size = text.size, hjust = 0, size.unit = "pt") +
  annotate("text", x = 2, y = 3.5, label = paste(Ki_0.90_0.80),
           size = text.size, hjust = 0, size.unit = "pt") +

  annotate("text", x = 1, y = 3.1, label = "85 - 95%",
           size = text.size, hjust = 0, size.unit = "pt") +
  annotate("text", x = 1, y = 2.9, label = paste(round(qnorm(0.85),2),"-", round(qnorm(0.95),2)),
           size = text.size, hjust = 0, size.unit = "pt") +
  annotate("text", x = 2, y = 3, label = paste(Ki_0.95_0.85),
           size = text.size, hjust = 0, size.unit = "pt") +

  annotate("text", x = 1, y = 2.6, label = "90 - 99.9%",
           size = text.size, hjust = 0, size.unit = "pt") +
  annotate("text", x = 1, y = 2.4, label = paste(round(qnorm(0.9),2),"-", round(qnorm(0.999),2)),
           size = text.size, hjust = 0, size.unit = "pt") +
  annotate("text", x = 2, y = 2.5, label = paste(Ki_0.999_0.90),
           size = text.size, hjust = 0, size.unit = "pt") +
```

```
  xlim(1,2.5) +
  ylim(0,5.2) +
  theme_void() +
  theme(plot.margin= margin(0,0,0,0))

p_2A.2 # show plot
```

| **Intervals of $K_i$ (% and probit)** | **Variability in $p_{50}$ (fraction of $\sigma$)** |
|---|---|
| 75 – 85%<br>0.67 – 1.04 | 0.36 |
| 80 – 90%<br>0.84 – 1.28 | 0.44 |
| 85 – 95%<br>1.04 – 1.64 | 0.61 |
| 90 – 99.9%<br>1.28 – 3.09 | 1.81 |

#### Illustrate the noise in p50 when Ki(%) restricted

##### Ki(%) restricted to 85-95%

```
# simulate 500 seed lots with random sigma-values (same as in (2) but
# initial viability restricted to range between 0.9 and 0.8)
lot_number <- 500 # number of random seed lots
viability <- runif(lot_number, min = 0.85, max = 0.95) # random initial viabilities for each
                                                        # seed lot varying between
                                                        # min (0.85) and max (0.95)
sigma <- runif(lot_number, min = 1, max = 50) # random values of sigma (in days) for each
                                               # seed lot  varying between
                                               # min (1 day) and max (50 days)
Simdat1 <- data.frame( # dataframe for simulated 500 random seed lots with
  viability = viability, # generated initial viabilities
  sigma = sigma) # and sigma
```

```
Simdat1$ID <- row(Simdat)[,1] # assign IDs to simulated species

# calculate p50 for simulated species
Simdat1 <- Simdat1 %>%
  mutate(p50 = qnorm(viability) * sigma)

p_2B <- ggplot(data = Simdat1, aes(x = sigma, y = p50, color = viability)) +
  geom_point(shape = 19, alpha = 0.6, size = point.size) +
  create_gradient(min(Simdat1$viability), max(Simdat1$viability)) +
  labs(x = "Sigma", y = expression(p[50])) +
  ggtitle(expression(K[i] ~"(%) = 85 - 95%")) +
  xlim(0,50) +
  ylim(0,150) +
  theme_classic() +
  theme(
    axis.title.x = element_text(size = lab.size),
    axis.title.y = element_text(size = lab.size),
    axis.text.x = element_text(size = text.size),
    axis.text.y = element_text(size = text.size),
    plot.margin = margin(c(0,0,0,0)),
    plot.title = element_text(size = lab.size))

p_2B # show plot
```

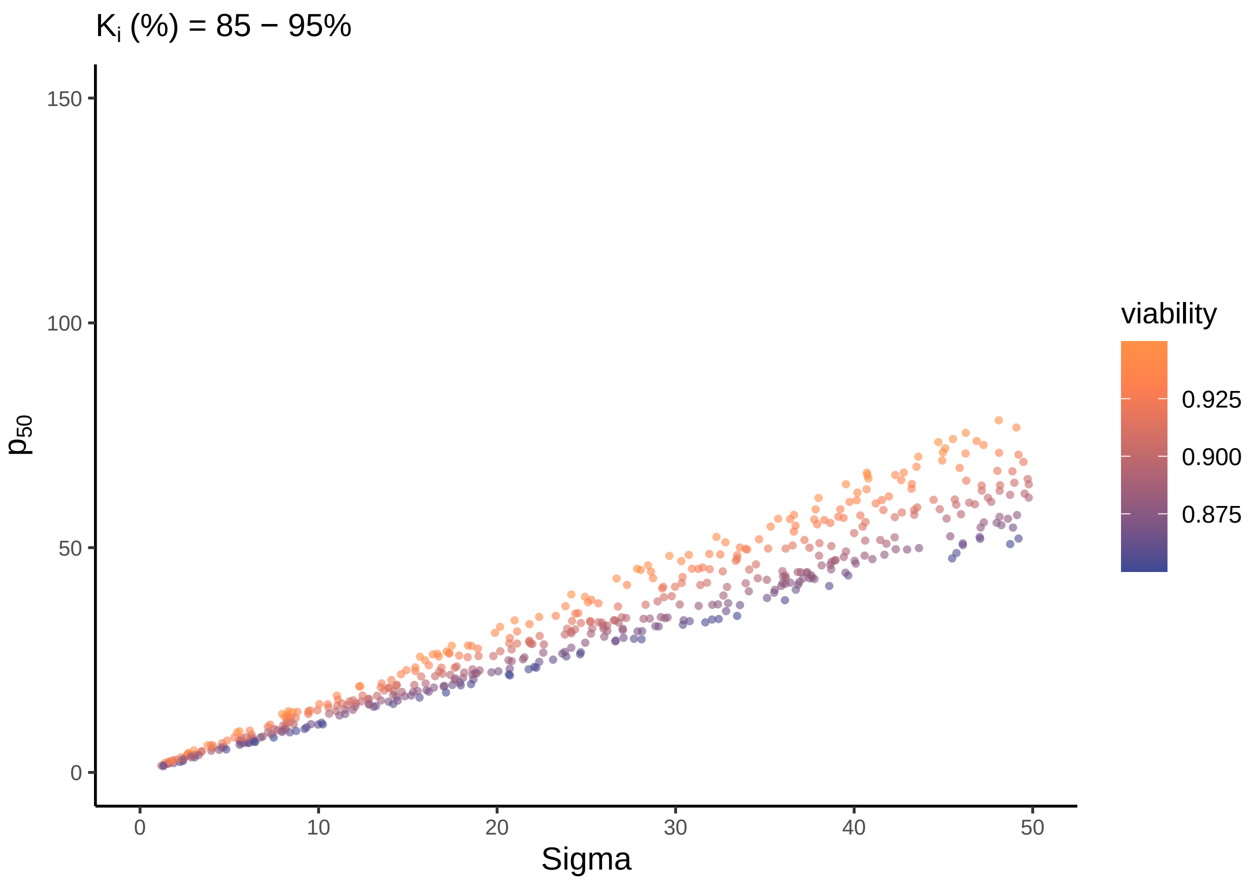

##### Ki(%) restricted to 90 to 99.9%

```
# simulate 500 seed lots with random sigma-values (same as in (2) but
# initial viability restricted to range between 0.999 and 0.90)
lot_number <- 500 # number of random seed lots
viability <- runif(lot_number, min = 0.90, max = 0.999) # random initial viabilities for each
                                                         # seed lot varying between
                                                         # min (0.90) and max (0.999)
sigma <- runif(lot_number, min = 1, max = 50) # random values of sigma (in days) for each
                                               # seed lot  varying between
                                               # min (1 day) and max (50 days)
Simdat2 <- data.frame( # dataframe for simulated 300 random seed lots with
  viability = viability, # generated initial viabilities
  sigma = sigma) # and sigma
Simdat2$ID <- row(Simdat)[,1] # assign IDs to simulated species

# calculate p50 for simulated species with original initial viability
Simdat2 <- Simdat2 %>%
  mutate(p50 = qnorm(viability) * sigma)  # Original p50 calculation

# Ki(%) limited to 0.9-0.999
p_2C <- ggplot(data = Simdat2, aes(x = sigma, y = p50, color = viability)) +
  geom_point(shape = 19, alpha = 0.6, size = point.size) +
  create_gradient(min(Simdat2$viability), max(Simdat2$viability)) +
  # scale_color_gradient2(low = "#374794", mid = "#966070", high = "#F57A50", midpoint =  0.85) +
  labs(x = "Sigma", y = expression(p[50])) +
  ggtitle(expression(K[i] ~"(%) = 90 - 99.9%")) +
  xlim(0,50) +
  ylim(0,150) +
  theme_classic() +
  theme(
    axis.title.x = element_text(size = lab.size),
    axis.title.y = element_text(size = lab.size),
    axis.text.x = element_text(size = text.size),
    axis.text.y = element_text(size = text.size),
    plot.margin = margin(c(0,0,0,0)),
    plot.title = element_text(size = lab.size))

p_2C # show plot
```

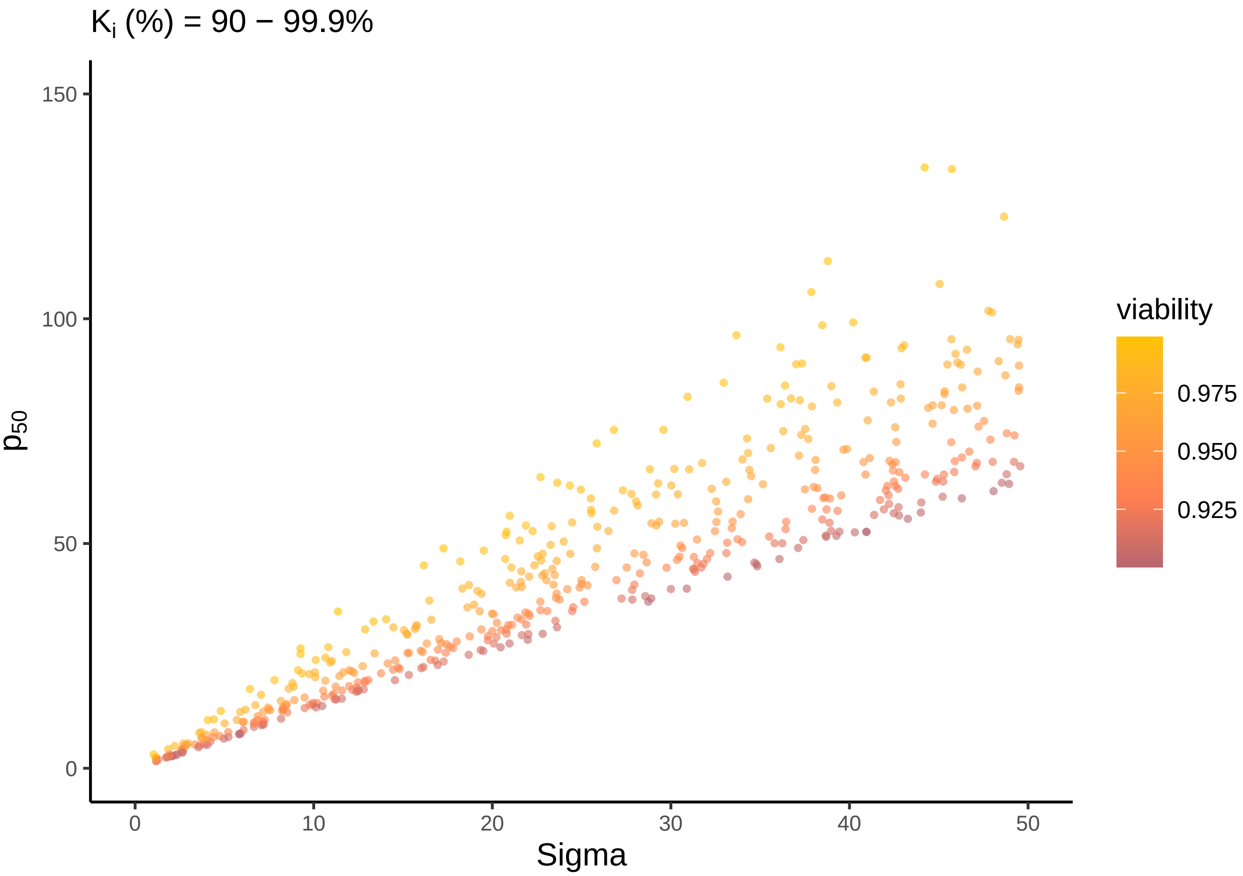


###### Combine Figure 2

```r
# remove legends from p_2B and p_2C
p_2B_nolegend <- p_2B + theme(legend.position = "none")
p_2C_nolegend <- p_2C + theme(legend.position = "none")

p_2 <- plot_grid(
  plot_grid(p_2A.1, p_2A.2, ncol = 2,
            labels = c("A", ""), label_size = 12, label_x = 0, label_y = 1,
            rel_widths = c(0.65, 0.35)),  # Adjust the width ratio for the first row
  plot_grid(p_2B_nolegend, p_2C_nolegend, p_legend_gradient, ncol = 3,
            labels = c("B", "C", ""), label_size = 12, label_x = 0, label_y = 1.01,
            rel_widths = c(0.42, 0.42, 0.16)),
  nrow = 2
) + theme(plot.margin = margin(5,15,0,0))

p_2 # show plot
```

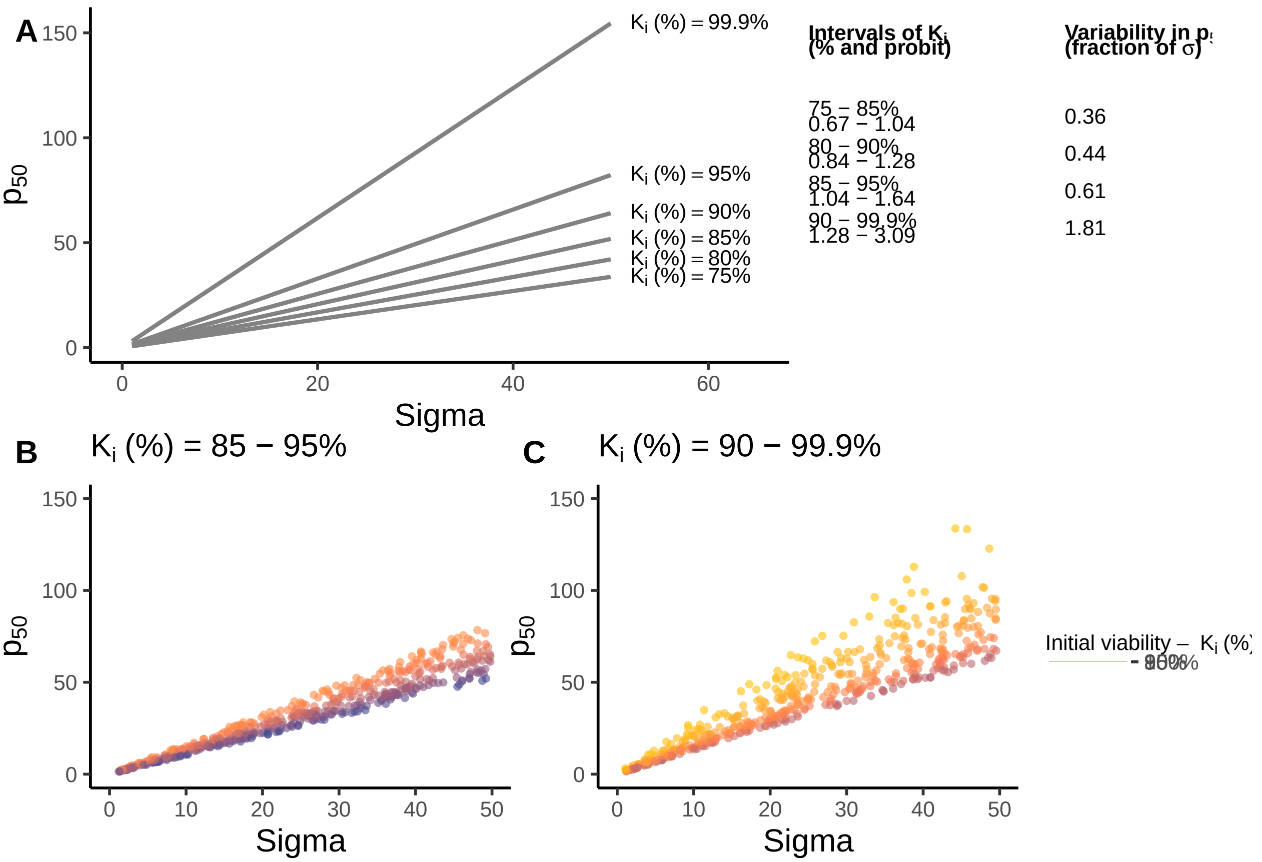


```
# save plot (the saved plots look identical to the Figures in the paper)
# ggsave(file = "your/path/Figure2.pdf", plot = p_2, width = 180, height = 200, units = "mm", dpi=500)
```

#### 3.2 Figure 3: Use sigma instead of p50

- Figure 3 helps to understand the parameter sigma.
- The code is rather long because it contains many design elements.

```
# simulate the extent of the viability loss in one sigma, depending on Ki, sigma is fixed to 1
Ki.percentage <- seq(from=0.999, to=0.023, by= -0.001) # generate initial viabilities
                                                        # between 2.3% and 99.9%
viab_sigma <- pnorm(qnorm(Ki.percentage)-1) # calculate viability after viability loss of one probit
                                            # (equals sigma days in ageing)
# qnorm(Ki.percentage): Probit transformation of initial viability,
# qnorm(Ki.percentage)- 1: substract 1 probit
# pnorm(qnorm(Ki.percentage)-1): Back transformation
loss <- Ki.percentage-viab_sigma # calculate viability loss in percentage points
                                 # after sigma days in ageing

# Create dataframe for viability loss depending on Ki
Viab_loss <- data.frame(
  Ki.percentage = Ki.percentage,
  viab_sigma = viab_sigma,
  loss = loss)
```

```r
# Set datapoints for three example initial viabilities to illustrate differences in viability loss
viab_A <- 0.691 # Example point A: 69.1% initial viability
viab_B <- 0.841 # Example point B: 84.1% initial viability
viab_C <- 0.977 # Example point C: 97.7% initial viability

# create dataframe with data for the 3 example points (A, B, C) --> necessary for figure 3
P_3_info <- data.frame(
  point = c("A", "B", "C"),
  viab_start = c(viab_A, viab_B, viab_C)
) %>%
  dplyr::mutate(
    viab_start_rounded = round(viab_start,2),
    viab_end = pnorm(qnorm(viab_start)-1),
    viab_start_time = (-1 * qnorm(viab_start) + 3),
    viab_end_time =  (-1 * qnorm(viab_end) + 3),
    viab_loss = viab_start-viab_end,
    text_y = (viab_start+viab_end)/2
  )

# generate subsets for each example point
P_3A <- P_3_info[P_3_info$point == "A",] # for p_3A
P_3B <- P_3_info[P_3_info$point == "B",] # for p_3B
P_3C <- P_3_info[P_3_info$point == "C",] # for p_3C

# Illustrate viability loss depending on Ki
p_3.1 <- ggplot(data = Viab_loss, aes(x = Ki.percentage, y = loss)) +
  geom_line(colour="#FF7F50") +
  geom_segment(aes(x = Inf, xend = P_3A$viab_start,
                   y = P_3A$viab_loss, xend = P_3A$viab_loss),
               color = "#374794", size = 0.6) +
  geom_segment(aes(x = Inf, xend = P_3B$viab_start,
                   y = P_3B$viab_loss, xend = P_3B$viab_loss),
               color = "#FFC300", size = 0.6) +
  geom_segment(aes(x = Inf, xend = P_3C$viab_start,
                   y = P_3C$viab_loss, xend = P_3C$viab_loss),
               color = "#9B4D96", size = 0.6) +
  geom_point(aes(x=P_3A$viab_start, y=P_3A$viab_loss), shape = 21,
             size = point.size+2.5, colour = "#374794", fill = "#B3C6E7") +
  geom_point(aes(x=P_3B$viab_start, y=P_3B$viab_loss), shape = 21,
             size = point.size+2.5, colour = "#FFC300", fill = "#FFF59D") +
  geom_point(aes(x=P_3C$viab_start, y=P_3C$viab_loss), shape = 21,
             size = point.size+2.5, colour = "#9B4D96", fill = "#E1BEE7") +

  # Text annotation
  annotate("text",
           x = P_3A$viab_start,
           y = P_3A$viab_loss,
           label =  bquote(K[i]~"(%)"~"="~.(P_3A$viab_start * 100)~"%"),
           hjust = -0.2,
           size = text.size,
           size.unit = "pt") +
  annotate("text",
           x = P_3B$viab_start,
```

```
           y = P_3B$viab_loss,
           label = bquote(K[i]~"(%)"~"="~.(P_3B$viab_start * 100)~"%"),
           hjust = -0.2,
           size = text.size,
           size.unit = "pt") +
  annotate("text",
           x = P_3C$viab_start,
           y = P_3C$viab_loss,
           label = bquote(K[i]~"(%)"~"="~.(P_3C$viab_start * 100)~"%"),
           hjust = -0.2,
           size = text.size,
           size.unit = "pt") +
  labs(x = expression("Initial viability - " ~ K[i] ~ " (%)"),
       y = "Viability loss per one sigma (% points)") +
  scale_x_reverse(labels = scales::percent) +
  scale_y_continuous(limits = c(0,0.4), labels = scales::percent) +
  # ylim(0.2,0.4) +
  theme_classic() +
  theme(
    axis.title.x = element_text(size = lab.size),
    axis.title.y = element_text(size = lab.size),
    axis.text.x = element_text(size = text.size),
    axis.text.y = element_text(size = text.size),
    plot.margin = margin(0,0,0,10))

p_3.1 # show plot
```

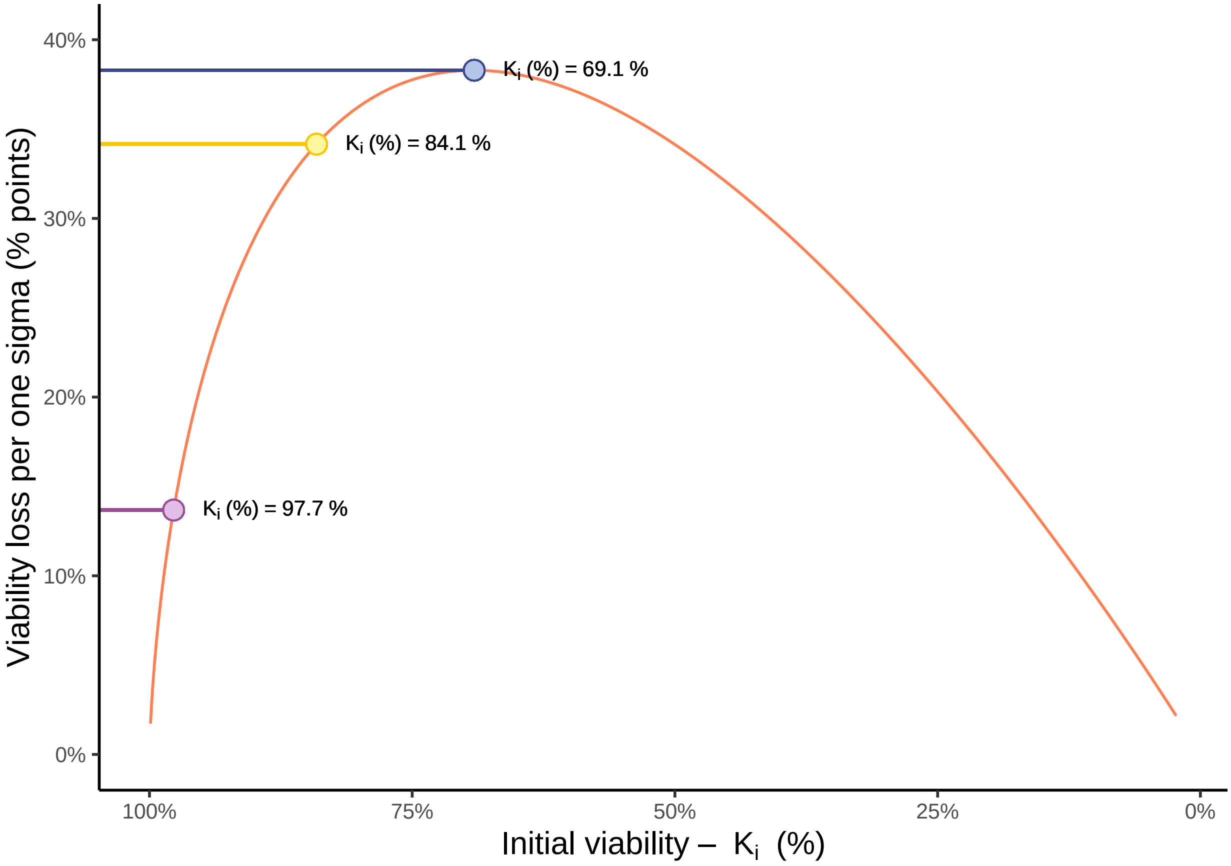


##### Illustrate viability loss for each of the three example points

Figure 3A

```
p_3A <- p_seed_survival +
  # horizontal line
  geom_segment(data = P_3A,
               aes(x = viab_start_time, xend = viab_end_time,
                   y = viab_end, yend = viab_end),
               color = "grey80", size = 0.6) +
  # Sigma annotation
  annotate("text",
           x = P_3A$viab_start_time + 0.5,
           y = P_3A$viab_end,
           hjust = 0.5,
           vjust = -0.3,
           label = expression(sigma), size = text.size, size.unit ="pt"
  ) +
  # Plot points from dataframe on seed survival
  geom_line(data = Seed_survival %>%
              dplyr::filter(seed_viability >= P_3A$viab_end & seed_viability <= P_3A$viab_start),
             size = line.size, colour = "#374794") +
  # Arrow
  geom_segment(data = P_3A,
               aes(x = viab_start_time, xend = viab_start_time,
```

```
                 y = viab_start, yend = viab_end),
               color = "#374794", size = 0.6,
               arrow = arrow(ends = "last", length = unit(0.02, "npc"))) +
  # Starting point viability loss
  geom_point(data = P_3A,
             aes(y = viab_start, x = viab_start_time),
             shape = 21, size = point.size + 2.5, colour = "#374794", fill = "#B3C6E7") +
  # Text: Viability loss
  annotate("text",
           x = P_3A$viab_start_time,
           y = 0.5 * (P_3A$viab_start + P_3A$viab_end),
           hjust = 1.1,
           label =
             paste0("Viability loss\n= ", round(P_3A$viab_loss, 3)*100, "% points"),
           size = text.size, size.unit ="pt") +
  ylab(expression("Initial viability - "~ K[i]~" (%)"))

p_3A # show plot
```

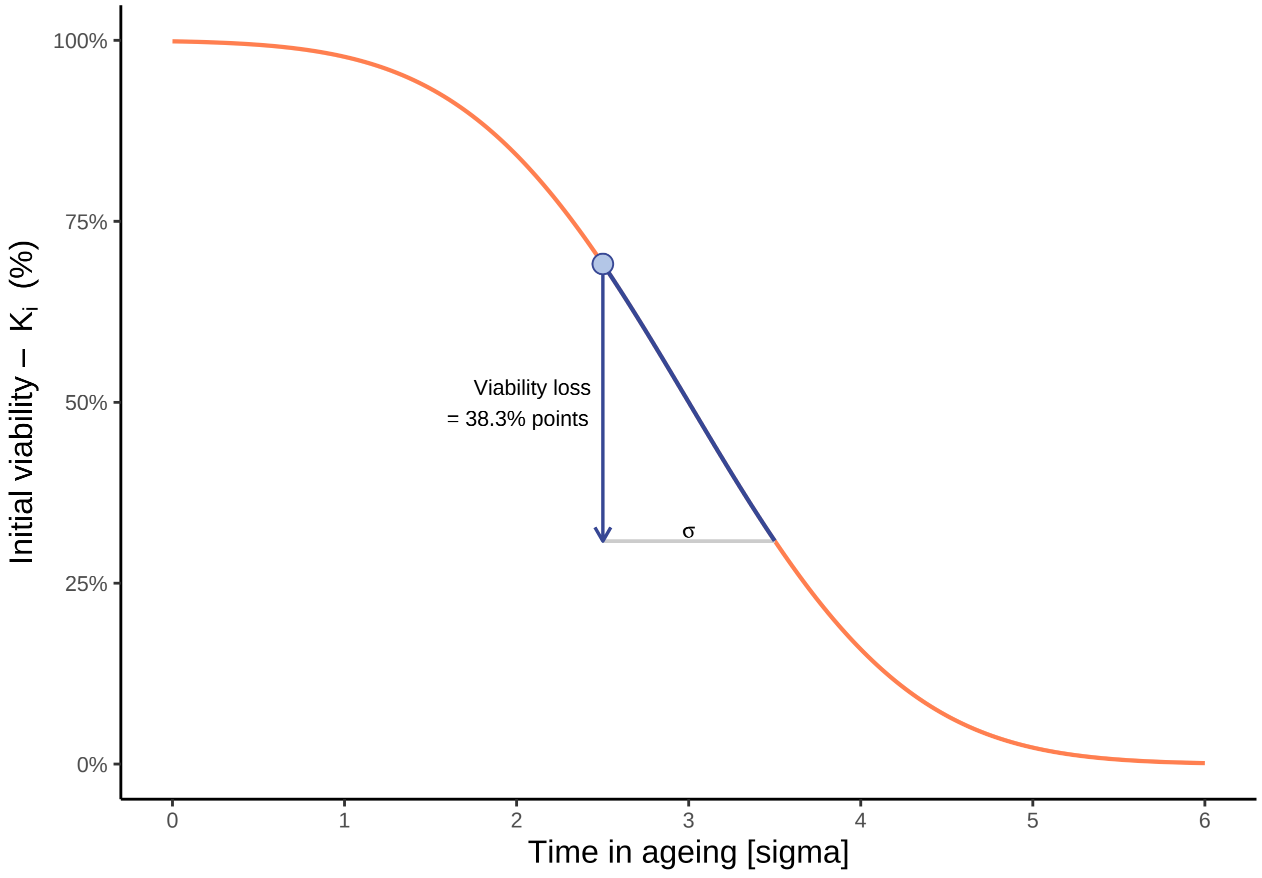


Figure 3 B

```
p_3B <- p_seed_survival +
  # horizontal line
  geom_segment(data = P_3B,
               aes(x = viab_start_time, xend = viab_end_time,
                   y = viab_end, yend = viab_end),
```

```r
                color = "grey80", size = 0.6) +
  # Sigma annotation
  annotate("text",
           x = P_3B$viab_start_time + 0.5,
           y = P_3B$viab_end,
           hjust = 0.5,
           vjust = -0.3,
           label = expression(sigma), size = text.size, size.unit ="pt"
  ) +
  # Plot points from dataframe on seed survival
  geom_line(data = Seed_survival %>%
              dplyr::filter(seed_viability >= P_3B$viab_end & seed_viability <= P_3B$viab_start),
            size = line.size, colour = "#FFC300") +
  # Arrow
  geom_segment(data = P_3B,
                aes(x = viab_start_time, xend = viab_start_time,
                    y = viab_start, yend = viab_end),
                color = "#FFC300", size = 0.6,
                arrow = arrow(ends = "last", length = unit(0.02, "npc"))) +
  # Starting point viability loss
  geom_point(data = P_3B,
              aes(y = viab_start, x = viab_start_time),
              shape = 21, size = point.size + 2.5, colour = "#FFC300", fill = "#FFF59D") +
  # Text: Viability loss
  annotate("text",
           x = P_3B$viab_start_time,
           y = 0.5 * (P_3B$viab_start + P_3B$viab_end),
           hjust = 1.1,
           label =
             paste0("Viability loss\n= ", round(P_3B$viab_loss, 3)*100, "% points"),
           size = text.size, size.unit ="pt") +
  ylab(expression("Initial viability - "~ K[i]~" (%)"))

p_3B # show plot
```

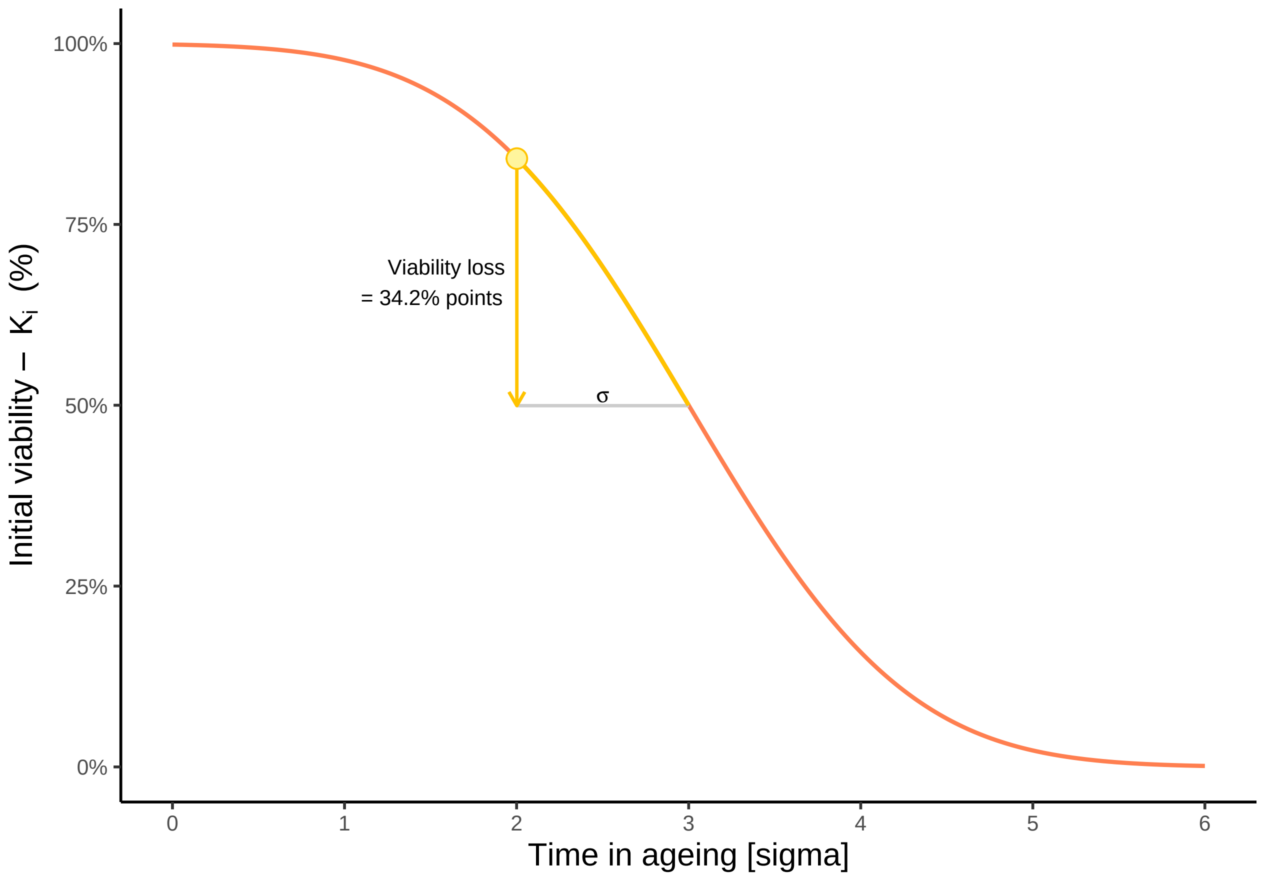


Figure 3C

```
p_3C <- p_seed_survival +
  # horizontal line
  geom_segment(data = P_3C,
               aes(x = viab_start_time, xend = viab_end_time,
                   y = viab_end, yend = viab_end),
               color = "grey80", size = 0.6) +
  # Sigma annotation
  annotate("text",
           x = P_3C$viab_start_time + 0.5,
           y = P_3C$viab_end,
           hjust = 0.5,
           vjust = -0.3,
           label = expression(sigma), size = text.size, size.unit ="pt"
  ) +
  # Plot points from dataframe on seed survival
  geom_line(data = Seed_survival %>%
              dplyr::filter(seed_viability >= P_3C$viab_end & seed_viability <= P_3C$viab_start),
            size = line.size, colour = "#9B4D96") +
  # Arrow
  geom_segment(data = P_3C,
               aes(x = viab_start_time, xend = viab_start_time,
                   y = viab_start, yend = viab_end),
               color = "#9B4D96", size = 0.6,
               arrow = arrow(ends = "last", length = unit(0.02, "npc"))) +
```

```
  # Starting point viability loss
  geom_point(data = P_3C,
             aes(y = viab_start, x = viab_start_time),
             shape = 21, size = point.size + 2.5, colour = "#9B4D96", fill = "#E1BEE7") +
  # Text: Viability loss
  annotate("text",
           x = P_3C$viab_start_time,
           y = 0.5 * (P_3C$viab_start + P_3C$viab_end),
           hjust = -0.45,
           label =
             paste0("Viability loss = ", round(P_3C$viab_loss, 3)*100, "% points"),
           size = text.size, size.unit ="pt") +
  ylab(expression("Initial viability - "~ K[i]~" (%)"))

p_3C # show plot
```

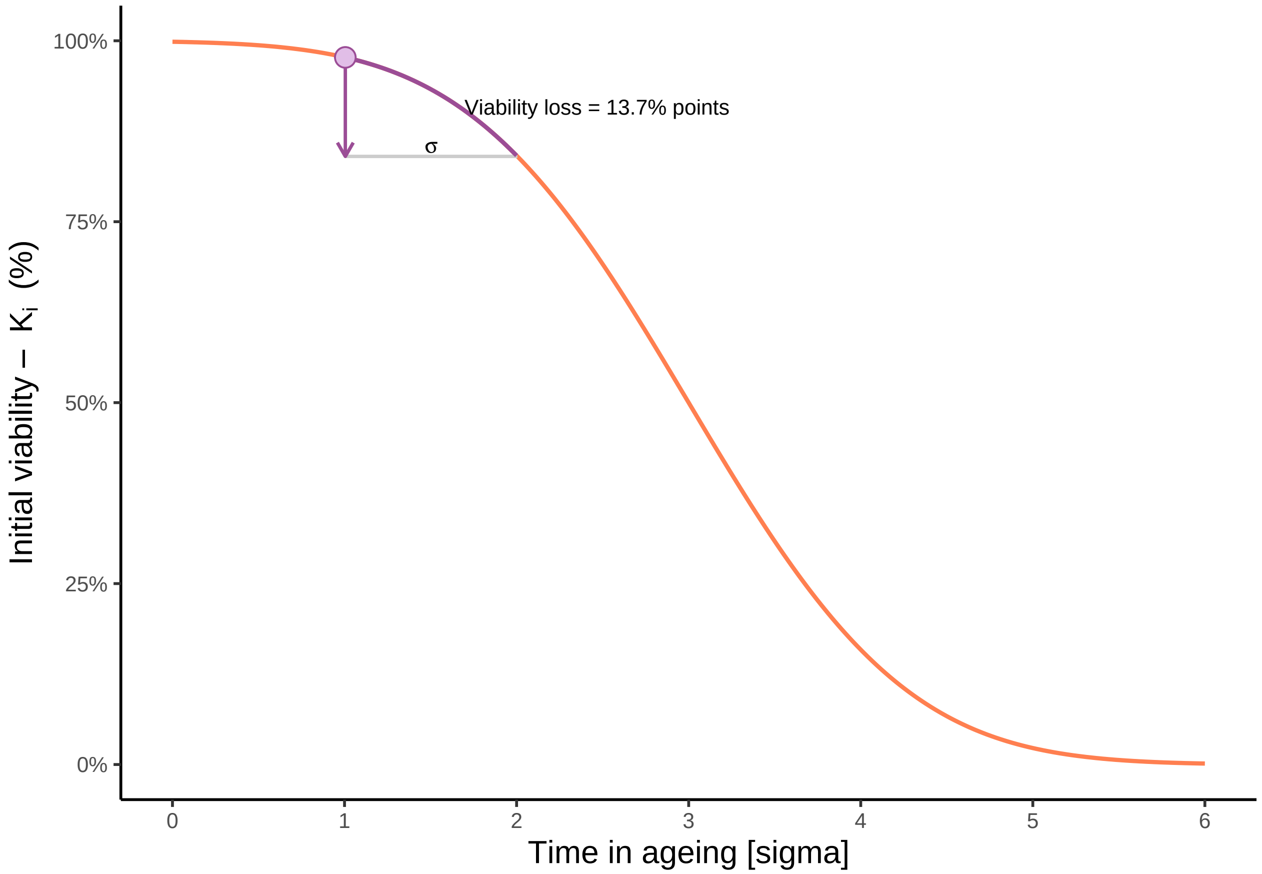


Combine plots for Figure 3

```
p_3 <- plot_grid(
  plot_grid(p_3A, p_3B, p_3C, ncol = 1, nrow = 3,
            labels = c("A", "B", "C"),
            label_size = 12, label_x = -0.01, label_y = 1),
  #           rel_heights = c(1, 1, 1)),
  p_3.1, ncol = 2,
  labels = c("", "D"), label_size = 12, label_x = 0, label_y = 1,
  rel_widths = c(0.5, 0.5)
```

```
)

p_3 # show plot
```

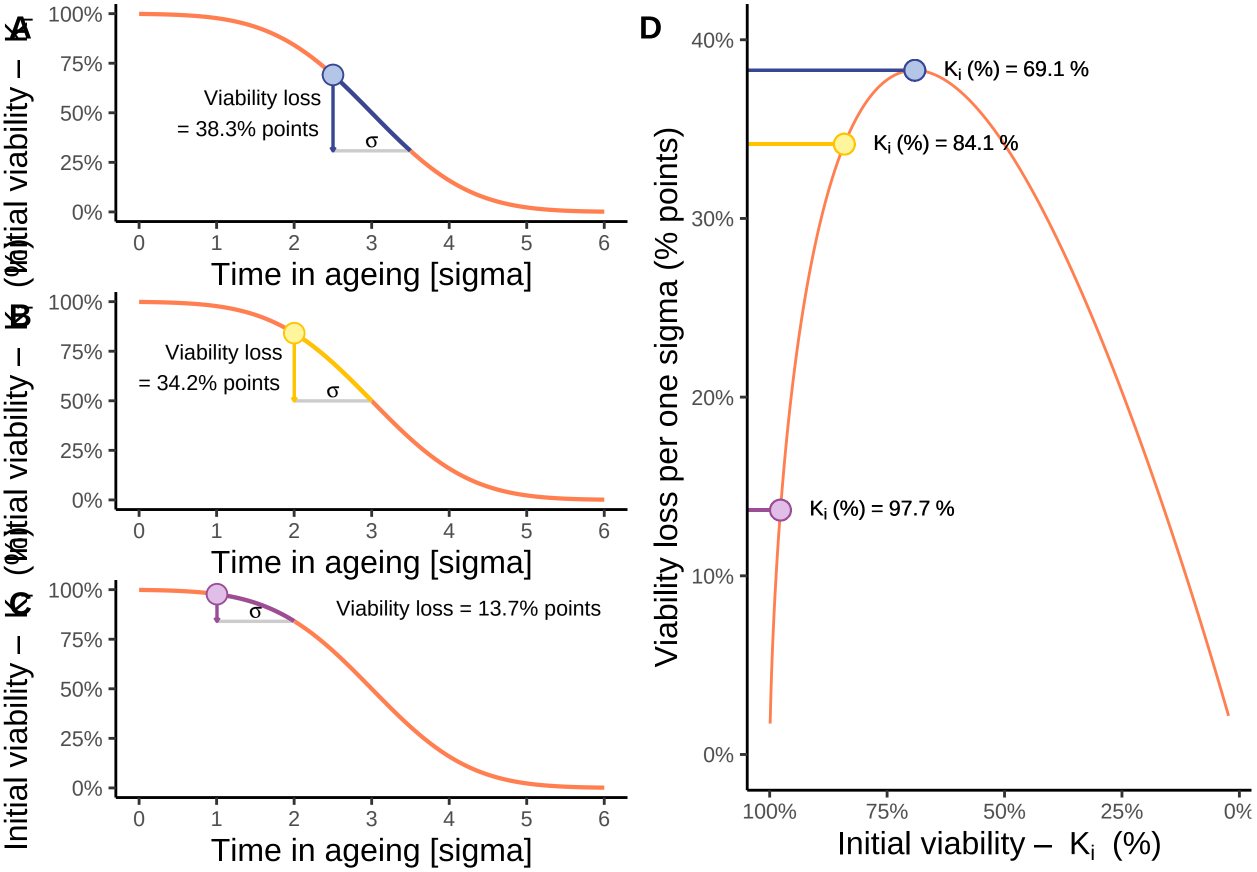


```
# save plot (the saved plots look identical to the Figures in the paper)
# ggsave(file= "your/path/Figure3.pdf", plot = p_3, width = 180, height = 200, units = "mm", dpi=500)
```

#### 3.3 Standardized p50

see Figure 2A
