## Supplementary Material 2 for "Limitations of *p*_50_ as a measure of seed longevity and the way forward"

### Limitations of p50 as a measure of seed longevity and the way forward (Calculations with 1 example seed lot)

Klepka, Lea
Bucharova, Anna

2025-05-16

#### Content:

0) Example data
1) Calculation of sigma, Ki and p50
2) Calculation of confidence intervals
3) Calculation of adjusted p50
4) Reciprocal calculation of seed longevity estimates

#### Load packages

```
library(ggplot2)
library(dplyr)
library(drc)
library(arm)
```

#### 0) EXAMPLE DATA

```
# Create vectors for each column
days_in_ageing <- c(0, 1, 5, 9, 15, 20, 30, 40, 57, 72)
sown <- rep(90, 10) # All 10 samples have 90 seeds
germinated <- c(88, 85, 86, 84, 84, 74, 42, 10, 1, 0) #
# Create the dataframe
Germination <- data.frame(days_in_ageing = days_in_ageing,
                          sown = sown,
                          germinated = germinated )
Germination # show dataframe
```

```
   days_in_ageing sown germinated
1               0   90         88
2               1   90         85
3               5   90         86
```

| | | | |
|---|---|---|---|
| 4 | 9 | 90 | 84 |
| 5 | 15 | 90 | 84 |
| 6 | 20 | 90 | 74 |
| 7 | 30 | 90 | 42 |
| 8 | 40 | 90 | 10 |
| 9 | 57 | 90 | 1 |
| 10 | 72 | 90 | 0 |

### (1) CALCULATION OF SIGMA, Ki AND p50

```r
# fit the seed survival model (probit analysis)
m <- glm(cbind(germinated, sown-germinated) ~ days_in_ageing,
         family = quasibinomial (link="probit"),
         data = Germination)
par(mfrow = c(2,2))
plot(m) # check model assumptions
```

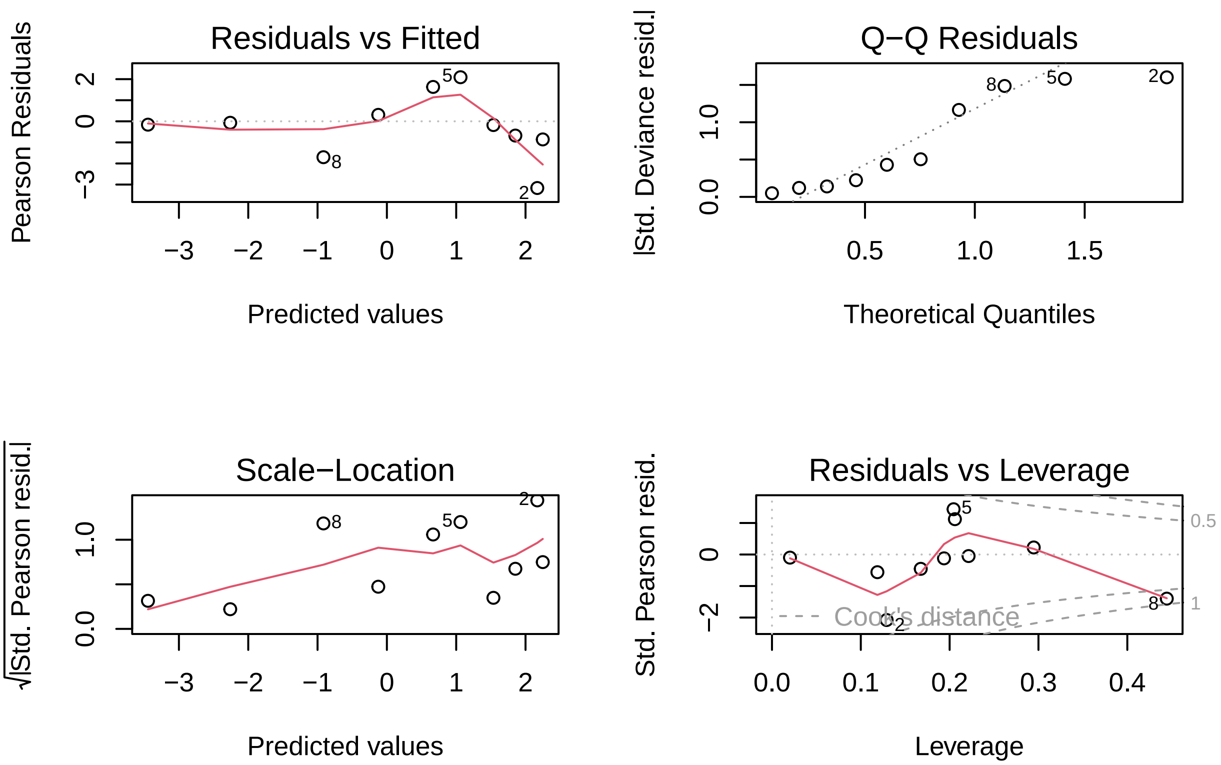


```r
coefs <- summary(m)$coefficients

Ki.probit <- coefs["(Intercept)", "Estimate"] # extract initial viability from model
                                              # on the probit scale
Ki.percentage <- pnorm(Ki.probit) # back-transformation of Ki (probit) to Ki(%)
```

```
sigma.probit <- coefs["days_in_ageing", "Estimate"] # extract slope (-1/sigma)
sigma.days <- -1/sigma.probit # get sigma value in days

p50.days <- Ki.probit*sigma.days # apply formula: p50 = Ki*sigma

# create summary table
t <- data.frame(Ki.probit, Ki.percentage, sigma.days, p50.days)
t # show table
```

```
  Ki.probit Ki.percentage sigma.days p50.days
1  2.248985     0.9877433   12.64156 28.43068
```

#### (2) CALCULATION OF CONFIDENCE INTERVALS

##### fit probit model as above (1)

```
m <- glm(cbind(germinated, sown-germinated) ~ days_in_ageing,
         family = quasibinomial (link="probit"),
         data = Germination)
par(mfrow = c(2,2))
plot(m) # check model assumptions
```

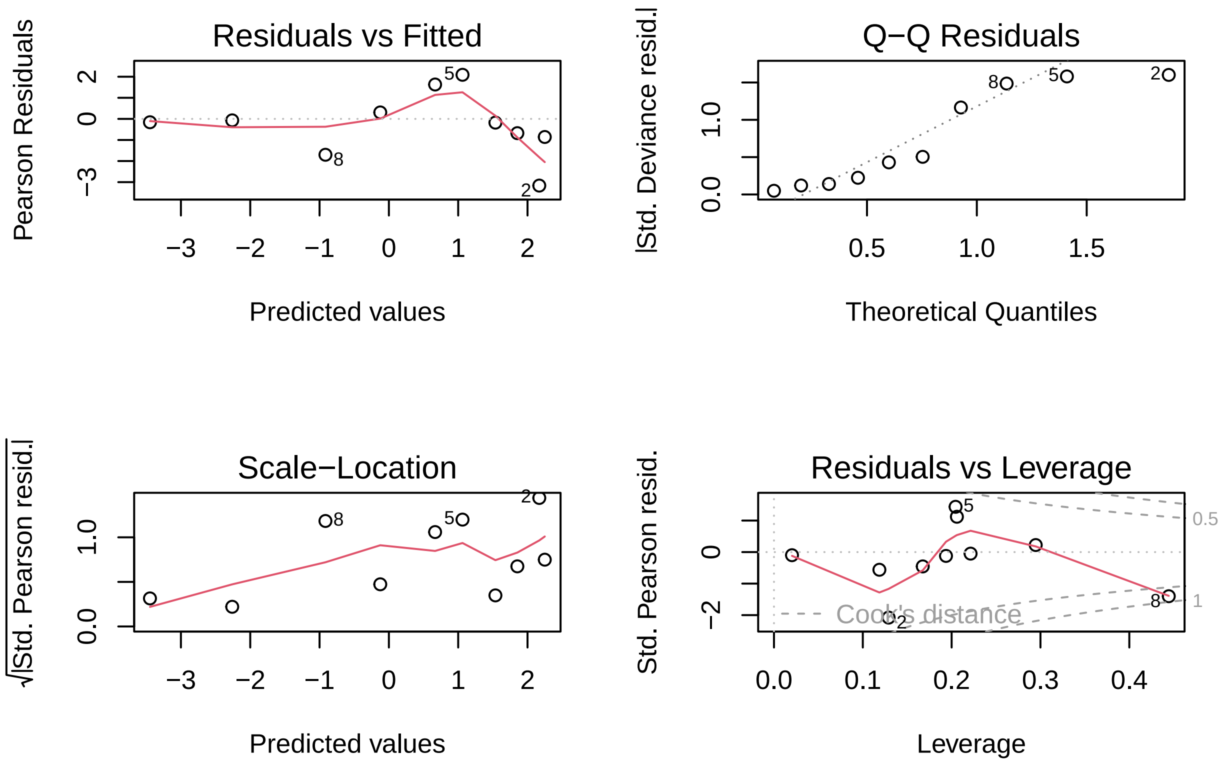

```
coefs <- summary(m)$coefficients
```

## (A) Ki

```
Ki.probit <- coefs["(Intercept)", "Estimate"] # extract initial viability from model
Ki.probit.SE <- coefs["(Intercept)", "Std. Error"] # extract Std. Error
Ki.probit.upper <- Ki.probit + 1.96*Ki.probit.SE # calculate upper limit of 95% CI
Ki.probit.lower <- Ki.probit -1.96*Ki.probit.SE # calculate lower limit of 95% CI
Ki.percentage.upper <- pnorm(Ki.probit.upper) # back transformation from probit scale
Ki.percentage.lower <- pnorm(Ki.probit.lower) # back transformation from probit scale
Ki.percentage <- pnorm(Ki.probit) # back-transformation of Ki (probit) to Ki(%)

# summary table for Ki
t.Ki <- data.frame(Ki.probit, Ki.probit.SE, Ki.probit.upper, Ki.probit.lower,
                   Ki.percentage.upper, Ki.percentage.lower, Ki.percentage)
t.Ki # show results
```

```
  Ki.probit Ki.probit.SE Ki.probit.upper Ki.probit.lower Ki.percentage.upper
1  2.248985    0.2048598         2.65051         1.84746           0.9959815
  Ki.percentage.lower Ki.percentage
1           0.9676597     0.9877433
```

#### (B) Sigma

```
sigma.probit <- coefs["days_in_ageing", "Estimate"] # extract sigma
sigma.probit.SE <- coefs["days_in_ageing", "Std. Error"] # extract Std. Error
sigma.probit.upper <- sigma.probit + 1.96*sigma.probit.SE # calculate upper limit of 95% CI
sigma.probit.lower <- sigma.probit - 1.96*sigma.probit.SE # calculate lower limit of 95% CI
sigma.days.upper <- -1/sigma.probit.upper # transformation to days
sigma.days.lower <- -1/sigma.probit.lower # transformation to days
sigma.days <- -1/sigma.probit #  transformation to days

# summary table for sigma
t.sigma <- data.frame(sigma.probit, sigma.probit.SE, sigma.probit.upper, sigma.probit.lower,
                      sigma.days.upper, sigma.days.lower, sigma.days)
t.sigma # show results
```

```
  sigma.probit sigma.probit.SE sigma.probit.upper sigma.probit.lower
1  -0.07910416     0.007714184        -0.06398436        -0.09422396
  sigma.days.upper sigma.days.lower sigma.days
1         15.62882         10.61301   12.64156
```

## (C) p50

We use Monte Carlo simulations for the confidence intervals of p50 because Ki and Sigma both have uncertainty around them (as Standard Error in the model) Simulation allows to avoid error multiplication

```
nsim <- 1000 # define number of simulations
bsim <- sim(m, nsim) # simulate nsim times the model m
apply(bsim@coef, 2, mean) # calculate mean for each column (Intercept and ageing duration)
```

```
   (Intercept) days_in_ageing
    2.24970139    -0.07924404
```

```
apply(bsim@coef, 2, quantile, prob = c(0.025, 0.975) ) # calculate 95% interval
```

```
      (Intercept) days_in_ageing
2.5%     1.845223    -0.09419641
97.5%    2.646205    -0.06366239
```

```
p50.vect <- bsim@coef[,1]*(-1/bsim@coef[,2]) # application of formula "p50 = Ki*sigma"
                                             # for each simulated model
p50.CI <- quantile(p50.vect, prob = c(0.025, 0.975)) # calculate 95% interval
                                                     # of all simulated p50 values
p50.upper <- as.numeric(p50.CI[2])
p50.lower <- as.numeric(p50.CI[1])

# summary table for p50
t.p50 <- data.frame(p50.days, p50.upper, p50.lower)
t.p50 # show results
```

```
  p50.days p50.upper p50.lower
1 28.43068  31.40903  25.39127
```

#### (3) CALCULATE ADJUSTED p50

```
Ki.std <- 0.9 # set this value as you like (in percentage)
p50.adj <- qnorm(Ki.std)*sigma.days # apply formula p50 = Ki*sigma (with sigma.days taken from (1))

# confidence intervals for adjusted p50
p50.adj.upper <- qnorm(Ki.std)*sigma.days.upper # calculate upper limit of 95% CI
                                                # (with sigma.days.upper taken from (2))
p50.adj.lower <- qnorm(Ki.std)*sigma.days.lower # calculate lower limit of 95% CI
                                                # (with sigma.days.lower taken from (2))

# summary table for p50
t.p50.adj <- data.frame(p50.adj, p50.adj.upper, p50.adj.lower)
t.p50.adj # show results
```

```
   p50.adj p50.adj.upper p50.adj.lower
1 16.20081      20.02914      13.60112
```

#### (4) RECIPROCAL CALCULATION OF SEED LONGEVITY ESTIMATES

Application of formula: p50 = Ki*sigma values for Ki, sigma (days) and p50 taken from (1)

#### (A) Calculate Ki from sigma and p50

```
Ki.probit <- p50.days/sigma.days # calculate Ki on probit scale
Ki.percentage <- pnorm(Ki.probit) # transform Ki from probit to percentage

Ki.probit # show result
```

```
[1] 2.248985
```

```
Ki.percentage # show result
```

```
[1] 0.9877433
```

#### (B) Calculate sigma from Ki and p50

```
Ki.probit <- qnorm(Ki.percentage)
sigma <- p50.days/Ki.probit

sigma # show result
```

```
[1] 12.64156
```

#### (C) Calculate p50 from sigma and Ki

```
Ki.probit <- qnorm(Ki.percentage)
p50.days <- Ki.probit*sigma.days

p50.days # show result
```

```
[1] 28.43068
```
